# Remodeling of perturbed chromatin can initiate *de novo* transcriptional and post-transcriptional silencing

**DOI:** 10.1101/2024.01.15.575522

**Authors:** Florian Carlier, Sebastian Castro Ramirez, Jaafar Kilani, Sara Chehboub, Isabelle Loïodice, Angela Taddei, Eugene Gladyshev

**Author notes:** Equal contribution (Florian Carlier, Sebastian Castro Ramirez).

## Abstract

In eukaryotes, repetitive DNA can become silenced *de novo*, either transcriptionally or post-transcriptionally, by processes independent of strong sequence-specific cues. The mechanistic nature of such processes remains poorly understood. We found that in the fungus *Neurospora crassa*, *de novo* initiation of both transcriptional and post-transcriptional silencing was linked to perturbed chromatin, which was produced experimentally by the aberrant activity of transcription factors at the *tetO* operator array. Transcriptional silencing was mediated by canonical constitutive heterochromatin. On the other hand, post-transcriptional silencing resembled repeat-induced quelling but occurred normally when homologous recombination was inactivated. All silencing of the *tetO* array was dependent on SAD-6 (a fungal ortholog of the SWI/SNF chromatin remodeler ATRX), which was required to maintain nucleosome occupancy at the perturbed locus. In addition, we found that two other types of sequences (the *lacO* array and native AT-rich DNA) could also undergo recombination-independent quelling associated with perturbed chromatin. These results suggested a model in which the *de novo* initiation of transcriptional and post-transcriptional silencing is coupled to the remodeling of perturbed chromatin.

**SIGNIFICANCE STATEMENT:** This study addresses an enigmatic question of how transcriptional and post-transcriptional gene silencing can be initiated *de novo* in the absence of strong sequence-specific cues. Using the fungus *Neurospora crassa* as a model organism, we found that both types of silencing can be triggered in mitotic cells by the remodeling of a transiently perturbed (nucleosome-depleted) chromatin state. In this system, the initiation of silencing requires SAD-6, a conserved SWI/SNF chromatin remodeler orthologous to ATRX that has been already implicated in repetitive DNA silencing in fungi, plants, and animals. Thus, the model proposed in this study may underpin a range of gene-silencing phenomena observed in other eukaryotes.

## INTRODUCTION

The genomes of most eukaryotes contain large amounts of repetitive DNA silenced transcriptionally and post-transcriptionally by diverse processes. While some of these processes are directed by strong sequence-specific signals (for example, KRAB domain-containing zinc finger proteins recruiting the methyltransferase SETDB1 [1]), others might be induced by the combined effect of multiple weaker interactions, necessitating a threshold number of tandem repeats to initiate silencing [2,3]. This principle may also apply to gene arrays (such as the human D4Z4 array carrying multiple copies of *DUX4* [4]) and transgenic repeats, as documented in fungi [5], plants [6], and animals [7,8]. Notably, in mammals, a screen for regulators of transgenic repeat-induced gene silencing (RIGS) identified a number of factors also required for transcriptional (heterochromatin-mediated) silencing of the native gene arrays [9,10], suggesting that the underlying mechanisms could be conserved.

ATRX (Alpha Thalassemia/Mental Retardation Syndrome X-Linked) is a member of the SWI/SNF family of chromatin remodelers [11]. In a complex with the histone chaperone DAXX, ATRX was shown to deposit the histone variant H3.3 at several types of repetitive sequences (pericentromeric, telomeric, ribosomal and other repeats) as well as non-repetitive regions [11]. H3.3 deposited by ATRX-DAXX often carries H3K9me3 [11]. Overall, ATRX has roles in several chromatin-based processes including transcriptional silencing of repetitive DNA, telomere maintenance, resolution of secondary DNA structures, and regulation of gene expression [11]. In addition to DAXX, ATRX is known to interact with other chromatin factors (proteins and long noncoding RNAs) as well as specific histone modifications (H3K9me3 and H3K4me0) [11]. ATRX is also known for its ability to recognize and remodel nucleosome-depleted chromatin linked to aberrant transcription, such as the Hsp70 genes during heat shock in Drosophila [12] and GC-rich telomeric repeats giving rise to R-loops [13].

The fungus *Neurospora crassa* sports several genome-defense processes that involve the *de novo* initiation of transcriptional and post-transcriptional silencing. First, in vegetative cells of *N. crassa*, transgenic repeats can become heavily methylated, as the result of potent RIGS [5]. Second, during sexual reproduction, duplications of genomic DNA longer than a few hundred base-pairs trigger repeat-induced point mutation (RIP), by which many cytosines in repeats are converted to thymines [14,15]. Interestingly, RIP can be carried out by the same pathway that establishes *de novo* DNA methylation during RIGS [5,16].

Concerning the initiation of post-transcriptional silencing, two processes need to be considered. One process, known as quelling, triggers the expression of small interfering RNAs (siRNAs) from repetitive transgenes in vegetative cells [17]. Another process, known as meiotic silencing by unpaired DNA (MSUD), takes place in early meiosis and induces the expression of siRNAs from the mismatching loci found at the allelic positions on pairs of homologous chromosomes [18,19]. Whereas MSUD occurs normally without DNA breakage and recombination [20], quelling was proposed to rely on recombinational DNA repair as a mechanism for repeat recognition [21]. Interestingly, potent MSUD was shown to require SAD-6, a fungal ortholog of ATRX [22].

A substantial part of the *N. crassa* genome corresponds to AT-rich DNA (produced by RIP), which nucleates H3K9me3 through a process mediated by the SUV39 methyltransferase DIM-5 [23]. In its turn, H3K9me3 is recognized by HP1 (Heterochromatin Protein 1), which forms a complex with the cytosine methyltransferase DIM-2 responsible for all DNA methylation in this organism [23]. Thus, in *N. crassa*, AT-rich DNA functions as a hard-wired signal for the formation of constitutive heterochromatin and ensuing DNA methylation [23]. Several additional components of this pathway were identified, including histone deacetylases and chromatin remodelers [24,25]. In parallel, a more dynamic type of DNA methylation occurs at some loci associated with antisense transcription [26,27]. Interestingly, this DNA methylation also requires DIM-5 and HP1 [28].

This study was motivated by two earlier results. First, in *N. crassa*, a long *tetO* operator array was reported to trigger RIP by the heterochromatin-related pathway, suggesting that it could initiate transcriptional silencing [29]. Second, in budding yeast, the association of such arrays with their corresponding repressor proteins was shown to induce transcriptional silencing of a nearby gene [30,31]. That process started with phosphorylation and concomitant depletion of histone H2A, suggesting that the binding of repressor proteins led to chromatin stress and nucleosome loss; and it did so by a mechanism independent of DNA replication [30]. The perturbed array was directed to the perinuclear compartment, where it became occupied by Silent Information Regulator (SIR) proteins, the main effectors of heterochromatic silencing in yeast [30]. Overall, it was proposed that the tight yet dynamic binding of repressor proteins could accelerate the turnover of nucleosomes and promote the incorporation of histones without certain modifications (such as H3K79me3), which would normally interfere with the recruitment of SIR proteins [30].

We now report that the *lacO* and the *tetO* operator arrays [32], when integrated as single-copy constructs and never exposed to RIP, can initiate strong transcriptional and post-transcriptional silencing in somatic cells of *N. crassa*. At the *tetO* array, the silencing was induced by the aberrant activities of either TetR-GFP or NIT-2 (a transcription factor of the GATA family). Both situations were associated with locally perturbed chromatin. Transcriptional silencing of this locus was mediated by canonical constitutive heterochromatin. On the other hand, post-transcriptional silencing resembled quelling but still occurred normally in the absence of RAD51 or RAD52, two main recombination factors. Therefore, this process was named recombination-independent quelling (RIQ). All silencing of the *tetO* array was dependent on SAD-6 (a fungal ortholog of the SWI/SNF chromatin remodeler ATRX), which was required to maintain nucleosome occupancy at the perturbed locus. In addition, we found that two other types of sequences (the *lacO* array and native AT-rich DNA) could also undergo recombination-independent quelling associated with perturbed chromatin. These results suggested a model in which the *de novo* initiation of transcriptional and post-transcriptional silencing is coupled to the remodeling of perturbed chromatin.

## RESULTS

### Experimental system

In budding yeast, the binding of bacterial repressor proteins to their corresponding operator arrays can trigger transcriptional silencing of a nearby reporter gene [31]. It was interesting to test if a similar process occurred in *N. crassa*, an organism with canonical transcriptional (heterochromatin-mediated) and post-transcriptional (RNAi-mediated) silencing [23]. To this end, the standard *lacO* and *tetO* arrays were used [32]. These arrays contained 168 *lacO1* and 191 *tetO* sites (interspersed with random sequences), and were characterized by GC-content of ∼40% and ∼50%, respectively (Fig. S1A). In addition, the *lacO* array included three perfect repeats of several hundred base-pairs (Fig. S1A). The arrays were concatenated on a plasmid and integrated between *his-3* and *lpl* by homologous recombination (Fig. 1A: ‘Strain A’). A synthetic construct expressing TetR-GFP was subsequently integrated as the replacement of *csr-1+* (Fig. 1A: ‘Strain B’). To protect the arrays from RIP and to eliminate the effects of cytosine methylation on H3K9me3 [23], the genes encoding DIM-2 and RID (a putative cytosine methyltransferase involved in RIP) were deleted in all strains used in this study. The state of chromatin was assayed by ChIP-seq and MNase-seq, while expression of small RNAs (sRNAs) was followed by sRNA-seq.

**Figure 1.**
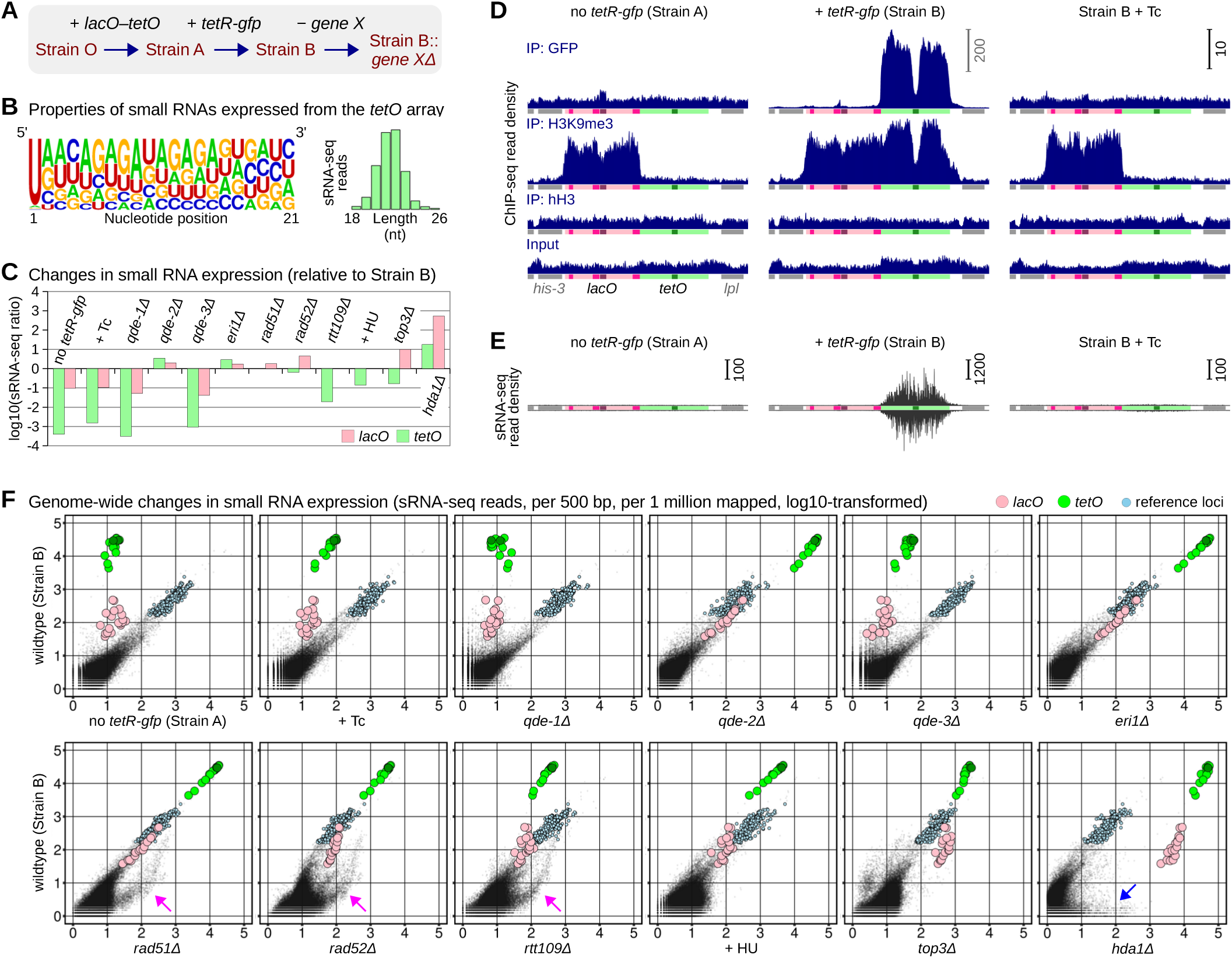
Repressor-bound *tetO* array initiates transcriptional and post-transcriptional silencing. (A) Series of strains created by homologous transformations, as indicated (strain identifications and complete genotypes are provided in Table S2). (B) Properties of sRNAs expressed from the repressor-bound *tetO* array (in Strain B). (C) Changes in the expression of array-derived sRNAs (see text for explanation). Corresponding raw data are plotted in Fig. 1F. (D) Density of ChIP-seq reads over the *lacO–tetO* reporter and the neighboring genes (per 1 bp, per 1 million mapped reads). Unless noted otherwise, all profiles are plotted to the same scale, corresponding to the default scale bar in the upper right corner. Tc: cultures were supplemented with tetracycline (100 μg/μl). The plotted region is 26000-bp long. (E) Density of sRNA-seq reads over the *lacO–tetO* reporter and the neighboring genes (analyzed and plotted as in Fig. 1D). (F) Scatter plots showing genome-wide changes in sRNA expression. The number of reads (expressed per 500 bp, per 1 million mapped reads) was augmented by 1 to enable log transformation. Tiles overlapping the *lacO* array, the *tetO* array, and the reference loci correspond to pink, green, and light-blue circles, respectively. Populations of sRNAs expressed from subtelomeric and AT-rich regions are marked by magenta and blue arrows, respectively.

### Binding of TetR-GFP to the *tetO* array induces strong transcriptional and post-transcriptional silencing

In the absence of TetR-GFP (in Strain A), only very low levels of sRNAs were expressed from the arrays (Fig. 1E,F; Fig. S1B). Interestingly, the *lacO* array was strongly enriched in H3K9me3 (Fig. 1D). An AT-rich motif ‘ATAACAATT’ was noted in the sequence of *lacO1* (Fig. S1A), raising a possibility that the *lacO* array could nucleate heterochromatin analogously to native AT-rich DNA in *N. crassa* [23].

Two remarkable results were observed at the *tetO* array in the presence of TetR-GFP. First, the array appeared strongly heterochromatic (Fig. 1D: ‘Strain B’). Second, it also started to express very large amounts of sRNAs (Fig. 1E,F: ‘Strain B’). These sRNAs were predominantly 20-23 nt long and had a strong bias for uracil at the 5’ position (Fig. 1B). Both processes were suppressed by tetracycline (Fig. 1D,E: ‘Strain B + Tc’), implicating the binding of TetR-GFP as a causative trigger of all silencing in this system.

Interestingly, the binding of TetR-GFP to the *tetO* array also stimulated sRNA expression at several loci as far as 100 kbp away from the array (Fig. S1B). The majority of those loci were non-coding. Such a long-distance effect may involve global repositioning of the array-carrying chromosomal segment, as reported previously in budding yeast [30], or increasing concentration of some critical RNAi factor in the vicinity of the array [33].

### A set of endogenous sRNA loci provides a standard reference for comparative sRNA-seq analysis

To compare expression of array-derived sRNAs between different conditions, a set of endogenous sRNA loci was selected as a standard reference. Such loci were chosen for their strong and invariant expression patterns, mostly corresponding to tRNA and 5S RNA genes (several examples are shown in Fig. S1C). For the purpose of this analysis, those loci were represented by 463 non-overlapping 500-bp tiles (SI Data File 3). For a given condition, the reference level was calculated as the median number of sRNA-seq reads mapped to those tiles. This value was used to normalize the levels of array-derived sRNAs calculated as the median number of reads mapped to the 500-bp tiles overlapping the arrays. Notably, this method could also be used to compare sRNA expression between *Neurospora* strains of different genetic origin (Fig. S1D).

### The *lacO–tetO* locus undergoes recombination-independent quelling (RIQ)

In *Neurospora*, expression of sRNAs from transgenic repeats represents a hallmark of quelling [17]. Thus, we asked if *tetO*-derived sRNAs required quelling factors QDE-1 (an RNA-dependent RNA polymerase), QDE-2 (an Argonaute protein), and QDE-3 (a RecQ helicase). The loss of QDE-1 suppressed *tetO*-derived sRNAs to the background levels corresponding to the repressor-free condition (Fig. 1C,F). The loss of QDE-3 produced a similar result, yet some residual expression of *tetO*-derived sRNAs above background could still be detected (Fig. 1C,F). In both conditions, the *tetO* array remained occupied by TetR-GFP (Fig. S2B). On the other hand, *tetO*-derived sRNAs did not require QDE-2, consistent with the fact that QDE-2 was dispensable for making sRNA during quelling (Fig. 1C,F). The levels of *lacO*-derived sRNAs, although being lower by two orders of magnitude compared to *tetO*-derived sRNAs, exhibited similar regulatory patterns (Fig. 1C,F).

We also tested the role of ERI1, a conserved exonuclease implicated in heterochromatin assembly and sRNA production associated with antisense transcription in *N. crassa* [34]. We found that ERI1 was not required for the expression of array-derived sRNAs in our system (Fig. 1C,F).

Earlier studies in *N. crassa* linked quelling with homologous recombination [21]. Thus, we asked if the newly uncovered quelling-like process required RAD51 and RAD52, the two critical recombination factors. Several effects were observed in the corresponding gene-deletion strains. First, the expression of *tetO*-derived sRNAs was not affected by the loss of RAD51 or RAD52 (Fig. 1C,F). Second, the expression of subtelomeric sRNAs became upregulated (Fig. 1F; Fig. S3C). Third, the proportion of background sRNAs (produced at low levels throughout the genome [35]) became strongly elevated as well (Fig. 1F; Fig. S3C). The loss of RAD52 had a greater impact compared to the loss of RAD51.

In *N. crassa*, the expression of some sRNA types requires the acetyltransferase RTT109 [36], which catalyses acetylation of H3K56 to control nucleosome dynamics coupled to DNA replication and repair [37]. We found that the loss of RTT109 had a strong impact on *tetO*-derived sRNAs, reducing their levels 52-fold (Fig. 1C,F). Interestingly, in this condition, the genome-wide pattern of sRNA expression resembled those observed in the absence of RAD51 or RAD52 (Fig. 1F; Fig. S3C). These results corroborated the previously reported role of RTT109 in sRNA biogenesis in *N. crassa* [36] and hinted at a possibility that RTT109 exerted its specific role in the production of *tetO*-derived sRNAs by modulating nucleosome homeostasis linked to DNA replication, rather than recombinational DNA repair.

Taken together, these results supported two conclusions. First, in *Neurospora,* similarly to budding yeast [31], the repressor-bound *tetO* array was engaged in transcriptional (heterochromatin-mediated) silencing. Second, the same locus was also engaged in post-transcriptional (RNAi-mediated) silencing, which resembled repeat-induced quelling but still occurred normally when homologous recombination was disabled. This process was named recombination-independent quelling (RIQ).

### Type I topoisomerase TOP3 is dispensable for RIQ

Sgs1 (the yeast ortholog of QDE-3) partners with the type I topoisomerase Top3 to form a conserved complex involved in recombinational DNA repair [38]. Thus, we asked if a *Neurospora* ortholog of Top3 played a role in RIQ. We found that the *top3Δ* condition was associated with a 6-fold decrease in *tetO*-derived sRNAs and a 10-fold increase in *lacO*-derived sRNAs (Fig. 1C,F). An elevated proportion of background sRNAs was noted as well (Fig. 1F). The latter effect is exemplified by having many sRNA-seq reads mapped to the active genes near the *lacO–tetO* locus (Fig. S3B). These results suggested that QDE-3 mediated RIQ by a mechanism that did not require TOP3.

### Heterochromatin assembly during RIQ is both RNAi- and recombination-independent

Our ChIP-seq analysis yielded several insights. First, the bulk levels of histone H3 at the *tetO* array decreased in the presence of TetR-GFP, indicating the occurrence of stressed (perturbed) chromatin (Fig. S2C). Second, the arrays remained perfectly heterochromatic when QDE-1, QDE-2, QDE-3, ERI1, RAD51, or RAD52 were removed individually (Fig. S2A,D), supporting the idea that, in *Neurospora,* constitutive heterochromatin and RNAi were largely independent from one another.

Interestingly, we found that the repressor-bound *tetO* array had its H3K9me3 levels decreased in the absence of RTT109 (Fig. S2A,D). This effect was not associated with apparent changes in the hH3 occupancy (relative to the parental strain), implying that TetR-GFP was still present normally (Fig. S2A,C). We used fluorescence microscopy as an independent approach to confirm this conclusion (Fig. S4). We note that the role of RTT109 in constitutive heterochromatin assembly was reported earlier in fission yeast [39].

### RIQ is constrained by constitutive heterochromatin

We next asked if the expression of array-derived sRNAs was constrained by constitutive heterochromatin. We found that the *lacO–tetO* locus lost all H3K9me3 in the absence of the histone deacetylase HDA1 (Fig. S2A). This process was linked to a decrease in the hH3 levels at the *lacO* array (Fig. S2C); and a comparable loss of hH3 also occurred at AT-rich DNA (Fig. S2C). With respect to sRNAs, two effects were observed. First, in the *hda1Δ* condition, the expression of *tetO*- and *lacO*-derived sRNAs surged 18- and 539-fold, respectively (Fig. 1C,F). Second, large amounts of sRNAs also became expressed from many AT-rich regions across the genome (Fig. 1F; Fig. S3A,C). A similar pattern was reported earlier in *Neurospora* strains lacking all heterochromatin [40].

### SAD-6 controls all silencing of the repressor-bound *tetO* array

An emerging connection between perturbed chromatin and RIQ encouraged us to investigate the involvement of chromatin remodeling factors. We started by testing CHD1 (Chromodomain Helicase DNA Binding Protein 1), which was already implicated in transcriptional and post-transcriptional silencing in *N. crassa* [21,26]. We found that the loss of CHD1 had no impact on the levels of hH3 or H3K9me3 at the arrays (Fig. S5A,C,D). To a first approximation, the levels of array-derived sRNAs were not affected as well (Fig. S5B).

Our results indicated that RIQ resembled MSUD in being recombination-independent. Therefore, we asked if the SWI/SNF chromatin remodeler SAD-6, which belongs to the ATRX clade and represents a critical MSUD factor [22], was involved in RIQ. Several surprising results were observed in the *sad-6Δ* condition. First, the expression of *tetO*-derived sRNAs decreased to a very low level associated with the absence of TetR-GFP and QDE-1 (Fig. 2A; Fig. S6A). We confirmed (by ChIP-seq and microscopy) that TetR-GFP was still present at the *tetO* array in this situation (Fig. 2B; Fig. S4). Second, the *tetO* array was no longer enriched in H3K9me3 (Fig. 2B). Third, the loss of silencing was accompanied by a strong decrease in the hH3 occupancy over DNA containing the *tetO* sites (Fig. 2B). Notably, hH3 was still retained at the *GmR* gene (in the middle of the *tetO* array); however, it was not associated with H3K9me3 (Fig. 2B). Fourth, the hH3 occupancy was restored in the presence of tetracycline (Fig. 2B). Fifth, similar dynamics was also exhibited by histone H2B (Fig. 2D).

**Figure 2.**
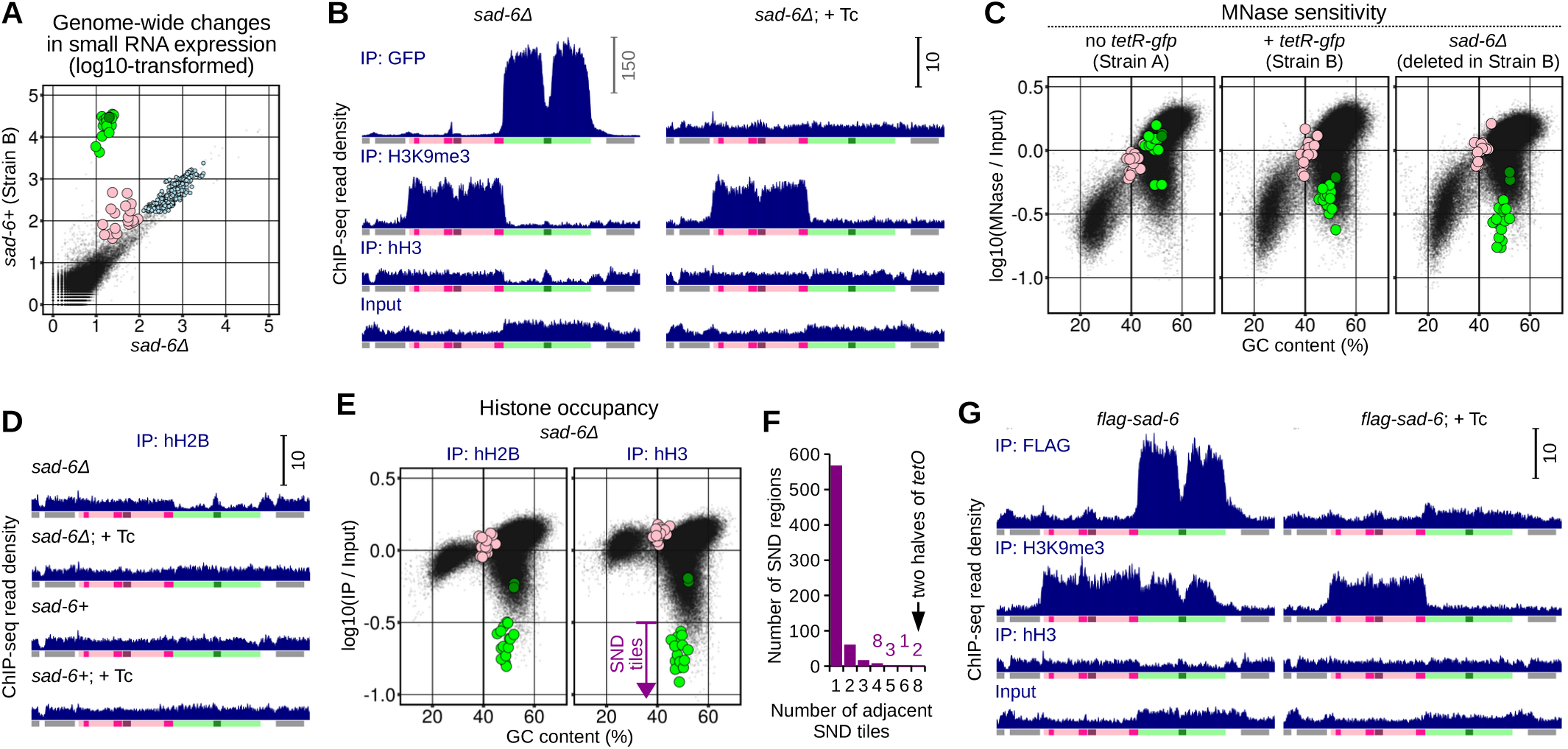
Chromatin remodeler SAD-6 controls all silencing of the repressor-bound *tetO* array. (A) Scatter plot showing genome-wide changes in sRNA expression (analyzed and plotted as in Fig. 1F). (B) Density of ChIP-seq reads over the arrays and the neighboring genes (analyzed and plotted as in Fig. 1D). (C) Scatter plots showing the relationship between MNase sensitivity and GC-content (plotted in Fig. S1E). Tiles overlapping the *lacO* and the *tetO* arrays are shown as pink and green circles, respectively. (D) Density of ChIP-seq reads over the arrays and the neighboring genes (analyzed and plotted as in Fig. 1D). (E) Scatter plots showing the relationship between histone occupancy and GC-content. Strongly nucleosome-depleted (SND) tiles were defined as those having the log10-transformed (IP[hH3]+1)/(Input+1) ratio of −0.5 or less. Tiles overlapping the *lacO* and the *tetO* arrays are shown as pink and green circles, respectively. (F) Histogram showing the relationship between the length and the number of strongly nucleosome-depleted (SND) regions. The length of SND regions corresponds to the number of consecutive SND tiles included in this region. (G) Density of ChIP-seq reads over the arrays and the neighboring genes (analyzed and plotted as in Fig. 1D).

The coordinated depletion of hH2B and hH3 (both encoded by single-copy genes in *N. crassa* [23]) pointed to the loss of entire nucleosomes. However, such an effect could also result from changes in epitope accessibility for ChIP. We used MNase sensitivity profiling to discriminate between these two possibilities. To this end, we found that the *tetO* array became progressively sensitive to MNase as its histone ChIP-seq coverage decreased (Fig. 2C,E; Fig. S2C), implying that the latter served as a reliable measure of nucleosome occupancy. Overall, the unperturbed *tetO* array was characterized by low and largely uniform MNase sensitivity (Fig. S7A: ‘Strain A’). Upon binding the repressor, the sensitivity of the array increased but still remained largely uniform (Fig. S7A: ‘Strain B’). This uniformity was disrupted in the absence of SAD-6 (Fig. S7A: *sad-6Δ*). Corresponding hH2B/hH3 ChIP-seq profiles hinted at a possibility that some strongly protected sites in the *sad-6Δ* condition could still feature nucleosomes in a fraction of the nuclei (Fig. S7B).

Genome-wide comparison of the *sad-6+*/*sad-6Δ* conditions yielded several results. First, the strong depletion of hH2B and hH3 in the *sad-6Δ* condition was restricted to the *tetO* array and rDNA (Fig. S6B,F). The latter result was intriguing, but it was not pursued in this study. Our further analysis revealed that the *tetO* array was the only genomic region incorporating more than 6 consecutive 500-bp ‘strongly nucleosome-depleted’ (SND) tiles (defined by having the log10-transformed (IP[hH3]+1)/(Input+1) ratio of −0.5 or less; Fig. 2E,F). Second, besides the *tetO* array, and possibly rDNA, constitutive heterochromatin was lost at only one additional locus, designated as ‘*r2*’ (Fig. S6B,F,J). Third, only one other unrelated locus, designated as ‘*r1*’, had its sRNA levels reduced in the absence of SAD-6 (Fig. S6E,I). In the wildtype, this region produced sRNAs from both strands by a process that also required QDE-1 (Fig. S3C; Fig. S6I). Fourth, the *tetO* array was the only genomic locus characterized by such a dramatic increase in MNase sensitivity (Fig. S6C,G).

### SAD-6 is specifically recruited to the repressor-bound *tetO* array

To test if SAD-6 regulated the *tetO* array directly, we replaced the promoter of the native *sad-6+* gene with a construct that (i) provided an N-terminal 3xFLAG tag and (ii) increased the expression of the tagged SAD-6 protein to compensate for a partial loss of its activity (Fig. S6D). We found that tagged SAD-6 became highly and specifically enriched at the repressor-bound *tetO* array, where it was partially active (as evidenced by the intermediate levels of hH3 and H3K9me3 at the array, Fig. 2G). Genome-wide, comparable enrichment was only observed at one additional locus (designated as ‘*r3*’), which overlapped the promoter of NCU01783 (Fig. S6H,K). The latter effect was unexpected yet reproducible (Fig. S6K). Taken together, these results suggested that the repressor-bound *tetO* array represented a particularly favorable substrate for SAD-6.

### NIT-2 is required for the *de novo* silencing of the *tetO* array induced by nitrogen starvation

Our results demonstrated that aberrant binding of TetR-GFP perturbed chromatin and initiated transcriptional and post-transcriptional silencing. It was critical to know if TetR-GFP played a general role in this process. If so, it could be replaced by an analogous unrelated protein. Here we took advantage of the fact that each *tetO* site contains two GATA motifs (Fig. S1A), which can recruit GATA factors *in vivo* [41]. In *Neurospora*, upon nitrogen starvation, the GATA factor NIT-2 is known to activate the expression of genes involved in nitrogen catabolism [42]. The DNA binding domain of NIT-2 is very conserved (Fig. 3A), and its capacity to recognize the GATA motif was demonstrated *in vitro* [42]. While only a few GATA motifs are found in the promoters of genes regulated by NIT-2 [43], the occurrence of many such motifs over the *tetO* array was expected to result in a situation where the aberrant activity of NIT-2 could perturb chromatin analogously to the strong binding of TetR-GFP.

**Figure 3.**
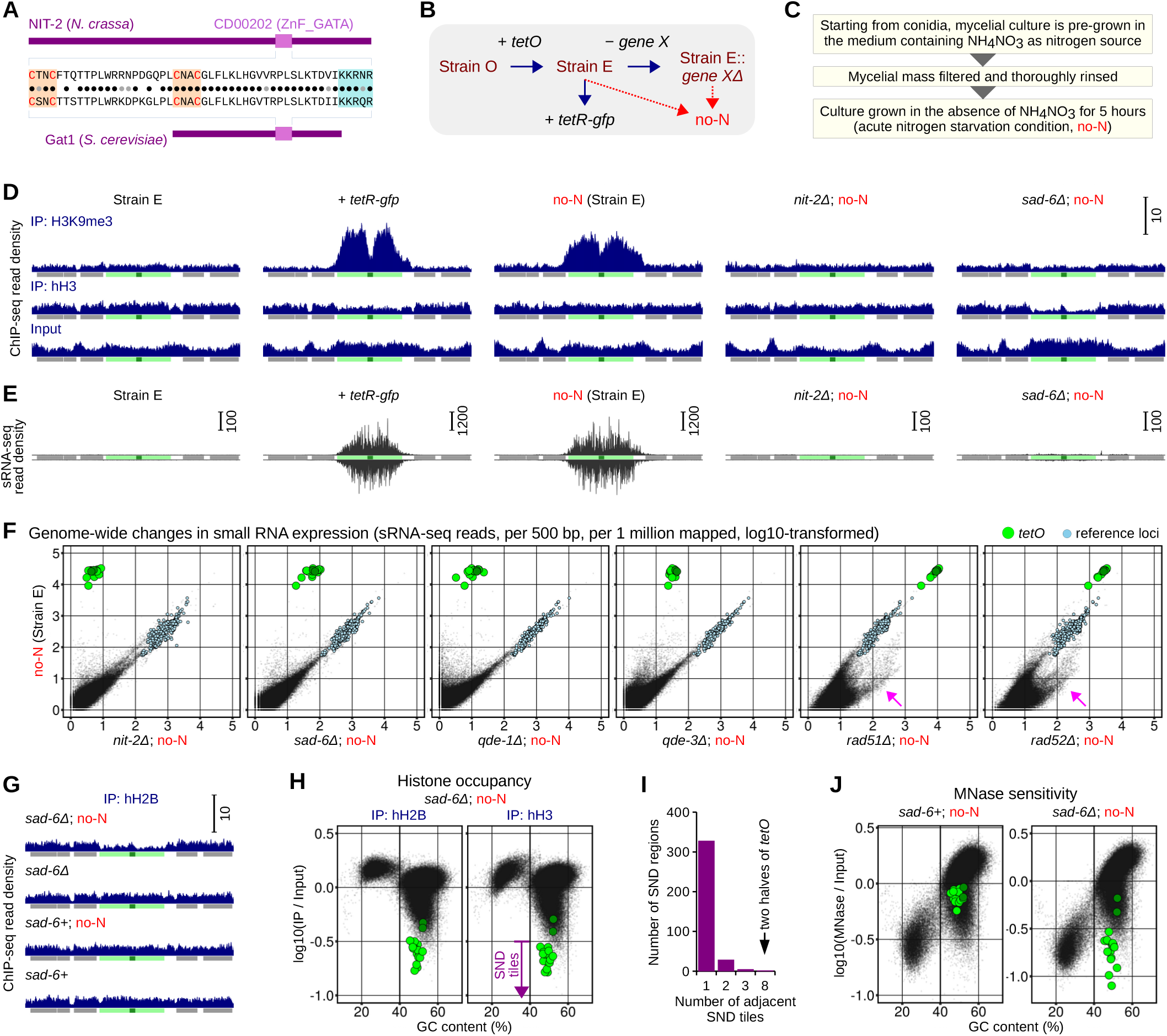
Nitrogen starvation induces silencing of the *tetO* array in the absence of the repressor. (A) Comparison of the DNA binding domains of NIT-2 and its yeast ortholog Gat1. (B) Series of strains created by homologous transformations, as indicated (strain identifications and complete genotypes are provided in Table S2). (C) Standard protocol for inducing acute nitrogen starvation. (D) Density of ChIP-seq reads over the array and the neighboring genes (analyzed and plotted as in Fig. 1D). (E) Density of sRNA-seq reads over the array and the neighboring genes (analyzed and plotted as in Fig. 1E). (F) Scatter plots showing genome-wide changes in sRNA expression (analyzed and plotted as in Fig. 1F). Tiles overlapping the *tetO* array and the reference loci are shown as green and light-blue circles, respectively. (G) Density of ChIP-seq reads over the array and the neighboring genes (analyzed and plotted as in Fig. 1D). (H) Scatter plots showing the relationship between histone occupancy and GC-content (nitrogen starvation in the absence of SAD-6; analyzed and plotted as in Fig. 2E). (I) Histogram showing the relationship between the length and the number of strongly nucleosome-depleted (SND) regions (analyzed and plotted as in Fig. 2F). (J) Scatter plots showing the relationship between MNase sensitivity and GC-content (analyzed and plotted as in Fig. 2C).

To test the role of NIT-2, we created a new strain carrying only the *tetO* array, without the adjacent *lacO* array (Fig. 3B: ‘Strain E’). This strain was also used to test for the *de novo* initiation of heterochromatin, as opposed to its spread from the *lacO* array. To this end, we found that, once TetR-GFP became available, the standalone *tetO* array expressed sRNAs and assembled H3K9me3 at the levels similar to those in Strain B (Fig. 3D,E: ‘*+ tetR-gfp*’), thus demonstrating the ability to initiate *de novo* transcriptional and post-transcriptional silencing.

Our basic nitrogen-starvation protocol included pre-growing *Neurospora* in the standard minimal medium for 24 hours, after which mycelial cultures were washed, transferred into a new medium lacking nitrogen, grown for additional 5 hours, and harvested for analyses (Fig. 3B,C). Thus, this protocol permitted assaying relevant parameters during the early stages of silencing. Remarkably, high levels of sRNAs and H3K9me3 were found at the *tetO* array upon nitrogen starvation, reaching those induced in the presence of the repressor (Fig. 3D,E). Critically, starvation-induced sRNAs and H3K9me3 were completely abrogated in the *nit-2Δ* condition (Fig. 3D,E,F). Genome-wide, several additional loci exhibited elevated levels of sRNAs upon starvation; for some of those loci this effect was dependent on NIT-2 (Fig. S8B).

Our further analysis revealed that the starvation-induced expression of *tetO*-derived sRNAs required QDE-1 and QDE-3 but not RAD51 or RAD52 (*tetO*-derived sRNAs decreased nearly 8-fold in the *rad52Δ* strain, yet they still exceeded background by two orders of magnitude; Fig. 3F). Thus, this process was classified as RIQ (Fig. 3F). Furthermore, starvation-induced assembly of constitutive heterochromatin occurred normally in the absence of QDE-1, QDE-3, RAD51 or RAD52 (Fig. S8A), excluding the roles of RNAi and recombination in its *de novo* initiation.

### SAD-6 controls all starvation-induced silencing

Thus far, our results indicated that the silencing triggered by nitrogen starvation (and mediated by NIT-2) was equivalent to the silencing induced by TetR-GFP. Subsequent experiments established that the corresponding *sad-6Δ* conditions were also equivalent (Fig. 3D-J). Specifically, the following effects were noted at the array upon nitrogen starvation in the *sad-6Δ* strain: *tetO*-derived sRNAs were not induced (Fig. 3E,F), constitutive heterochromatin was not assembled (Fig. 3D), and the array became strongly nucleosome-depleted as well as MNase-hypersensitive (Fig. 3D,G-J). Interestingly, the array was essentially the only genomic locus featuring the aforementioned chromatin defects (Fig. S8C,D).

Comparison of high-resolution MNase-seq profiles yielded additional insights. First, MNase sensitivity of the unperturbed *tetO* array was not influenced by the adjacent heterochromatic *lacO* array (Fig. S7A,C: ‘Strain A’ and ‘Strain E’). Second, when Strain E was starved for nitrogen, MNase sensitivity of the *tetO* array increased but remained largely uniform (Fig. S7A,C). Third, this uniformity was compromised in the *sad-6Δ* condition, recapitulating the state of the *tetO* array bound by TetR-GFP in the absence of SAD-6 (Fig. S7A,C). Notably, the two *sad-6Δ* conditions featured MNase-seq profiles that were markedly different, suggesting that the fine structure of the perturbed chromatin state was influenced by the nature of the perturbing agent.

### DNA replication is required for starvation-induced RIQ

Canonical repeat-induced quelling can be suppressed by 0.1M hydroxyurea (HU) [44], a potent yet reversible inhibitor of DNA replication in *N. crassa* [45]. We also found that strong RIQ of the *tetO* array was dependent on the acetyltransferase RTT109 with known roles in DNA replication (Fig. 1C,F) [37]. Therefore, we asked if DNA replication was required for RIQ. Focusing first on the repressor-bound *tetO* array, we discovered that a 24-hour incubation of pre-grown mycelia in the minimal medium containing 0.1M HU downregulated *tetO*-derived sRNAs by a factor of 7 (while having no effect on *lacO*-derived sRNAs; Fig. 1C,F: ‘Strain B + HU’). By microscopy, we confirmed that the repressor was still localized properly upon the HU treatment (Fig. S4). The enlarged size of the HU-treated nuclei and the increased proportion of background sRNAs were noticed (Fig. 1F; Fig. S4). This result indicated a possible connection between RIQ and DNA replication, yet because HU was added to the pre-grown mycelial cultures with already active RIQ, the magnitude of this effect could not be determined with certainty.

The above question was addressed using an experimental system provided by starvation-induced RIQ (Fig. 4). To block all DNA replication before starvation, HU was added during the last 5 hours of normal growth (Fig. 4A). Subsequently, the HU block could be either released or maintained during the starvation phase (Fig. 4A). Using this system, we found that maintaining the HU block suppressed all starvation-induced RIQ of the *tetO* array (Fig. 4B,C). The only other co-suppressed sRNAs were those expressed from AT-rich DNA (Fig. 4C).

**Figure 4.**
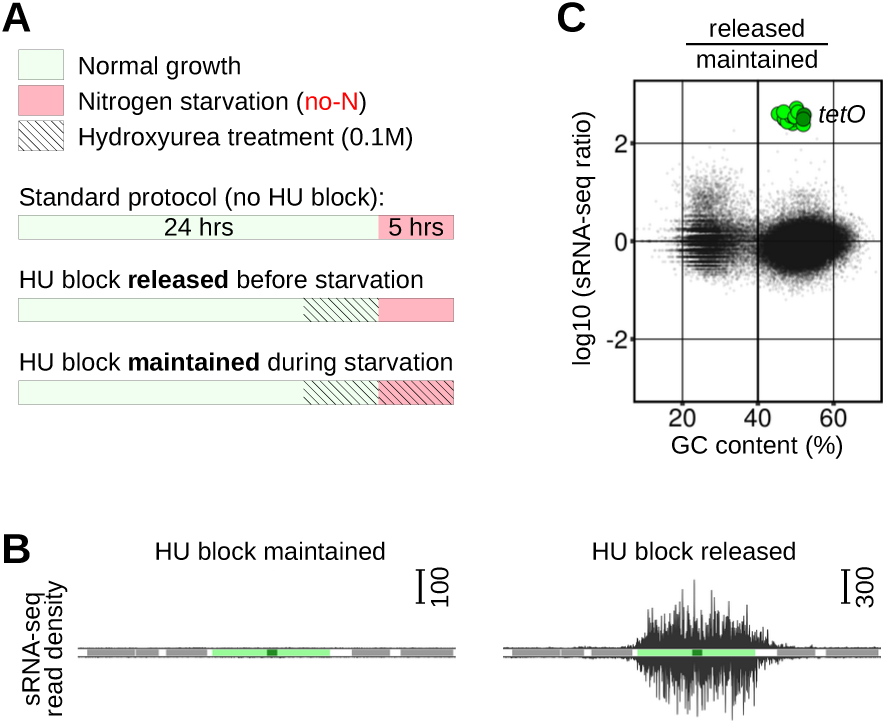
DNA replication is required for post-transcriptional silencing induced by nitrogen starvation. (A) Modifications of the standard nitrogen-starvation protocol (Fig. 3C), which include treatments with 0.1M hydroxyurea (HU) before and (optionally) during starvation. (B) Density of sRNA-seq reads over the array and the neighboring genes (analyzed and plotted as in Fig. 1E). (C) Scatter plot showing the relationship between changes in sRNA expression and GC-content.

### The *lacO* array and native AT-rich DNA can undergo RIQ

Our results suggested that native AT-rich DNA and the *lacO* array shared the capacity to nucleate constitutive heterochromatin. In addition to constitutive heterochromatin, *Neurospora* features facultative heterochromatin marked by H3K27me3, which is deposited by the lysine methyltransferase SET-7 [23]. Compromised fitness of *dim-5Δ* strains can be partially restored by deleting *set-7+* [40,46]. Therefore, to analyze the expression of sRNAs from the *lacO* array and AT-rich DNA, we moved the *lacO–tetO* reporter to a new genetic background in which *dim-5+* and *set-7+* were both deleted (Fig. 5A: ‘Strain C’). A matching *dim-5+* strain was created by replacing *csr-1+* with the *dim-5+* transcription unit [16] (Fig. 5A: ‘Strain D’). Our analysis of these otherwise isogenic strains suggested that DIM-5 promoted hH3 occupancy over AT-rich DNA and the *lacO* array, while also suppressing sRNA expression from these same regions (Fig. 5B,C; Fig. S9A). Affected sRNAs required QDE-1 but not RAD51, and they were also downregulated by HU (Fig. 5D; Fig. S9B-D). The roles of QDE-3 and RAD52 were not tested, because we were unable to delete *qde-3+* or *rad52+* in our basic *dim-5Δ* strain.

**Figure 5.**
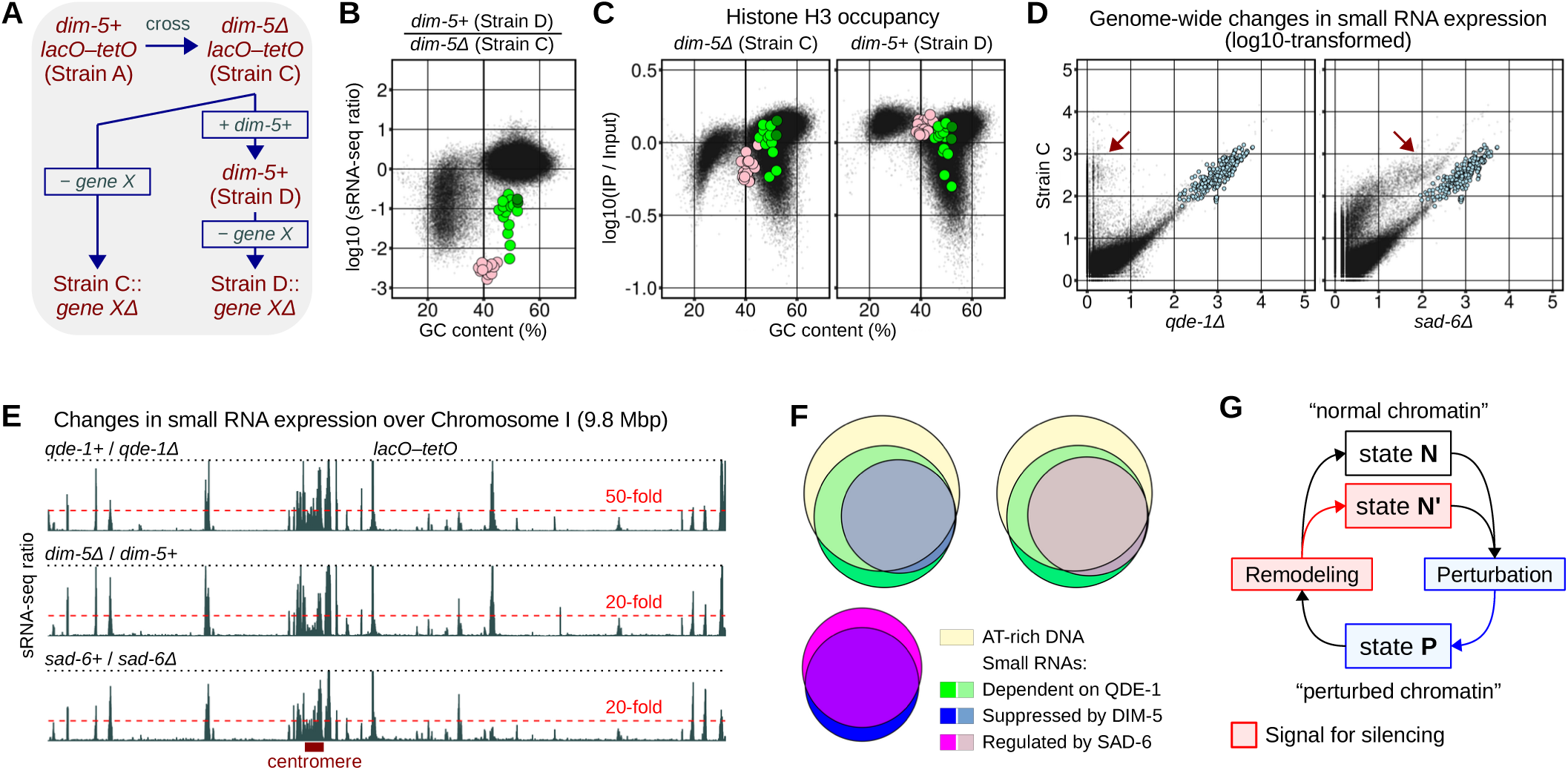
SAD-6 also regulates post-transcriptional silencing of native AT-rich DNA. (A) Series of strains created by homologous transformations, starting from Strain C (strain identifications and complete genotypes are provided in Table S2). Strain C was produced by crossing Strain A with another strain carrying *dim-5Δ* and *set-7Δ* (Table S2). (B) Scatter plot showing the relationship between changes in sRNA expression and GC-content (analyzed and plotted as in Fig. 4C). Tiles overlapping the *lacO* and the *tetO* arrays are shown as pink and green circles, respectively. (C) Scatter plots showing the relationship between histone occupancy and GC-content (analyzed and plotted as in Fig. 2E). (D) Scatter plots showing genome-wide changes in sRNA expression (analyzed as in Fig. 1F). Populations of sRNAs affected differentially by the deletions of *qde-1+* and *sad-6+* in Strain C are indicated (brown arrows). Reference loci are shown as light-blue circles. (E) Changes in sRNA expression across Chromosome I, obtained as ratios of sRNA-seq reads using the same array of 500-bp tiles as in Fig. 1F. All denominator values were augmented by 1. (F) Euler diagrams showing the relationships among several groups of genomic loci (represented by the same set of 500-bp tiles as in Fig. 1F). AT-rich DNA is defined by GC-content ≤ 38%. Other thresholds correspond to the five-fold change in sRNA expression. (G) Chromatin can exist in two states, normal and perturbed. Perturbed state can be promoted by the aberrant activities of DNA-binding proteins, such as TetR-GFP or NIT-2. Chromatin can be returned to its normal state by dedicated chromatin remodelers, such as SAD-6, which are also required to initiate the *de novo* silencing.

In the *dim-5Δ* condition, two results concerning the function of SAD-6 at AT-rich DNA were observed. First, SAD-6 was required for the optimal expression of most AT-rich sRNA loci suppressed by DIM-5 (Fig. 5D-F). Second, the loss of SAD-6 was associated with a broad increase in MNase sensitivity of AT-rich DNA (Fig. S9E,F). This effect was not evident for heterochromatinized AT-rich DNA (Fig. S6C). For the majority of the AT-rich loci engaged in RIQ, the increase in MNase sensitivity was moderate and largely uniform (Fig. S9F).

### The *lacO* array becomes genetically unstable in the absence of heterochromatin

We found that the *lacO* array could still undergo RIQ in the absence of SAD-6 (Fig. S9D). This result was not surprising, as SAD-6 was also partially dispensable for RIQ of AT-rich DNA. However, in this *sad-6Δ* strain, approximately one half of the *lacO* array was deleted (Fig. S9D). Because such deletions were never observed in the other *sad-6Δ* strains, this result indicated that the *lacO* array became genetically unstable when stripped of heterochromatin. We used multiplex PCR to test the stability of the arrays in clonal populations of nuclei of several basic strains (Fig. S10). Two pairs of primers were mixed for each PCR. One pair was specific for the central portion of either the *lacO* or the *tetO* array, the second pair amplified a portion of *spo11* as the positive control (Fig. S10B). Using this approach, frequent deletions in the *lacO* array were uncovered, but only in the heterochromatin-deficient background (Fig. S10B: ‘Strain C’). Such deletions corresponded to recombination products involving the three perfect repeats present in the *lacO* array (Fig. S1A). Thus, the *lacO* array differed from the *tetO* array in becoming genetically unstable in its perturbed state.

### HDA1 and CHAP, members of the HCHC complex, play different roles at the *lacO* array

We next analyzed the role of constitutive heterochromatin in RIQ of the *lacO* array. In *N. crassa*, constitutive heterochromatin is controlled by several protein complexes, most notably by DCDC and HCHC [23]. DCDC catalyzes H3K9me3 and contains DIM-5 (the catalytic subunit), DIM-7, DIM-9, CUL4, and DDB1; whereas HCHC mediates histones deacetylation and contains HDA1 (the catalytic subunit), along with HP1, CDP-2, and CHAP [23]. With respect to DCDC, the loss of DIM-7 or DIM-9 affected the *lacO* array similarly to the loss of DIM-5: H3K9me3 was absent, the hH3 occupancy decreased, and RIQ became strongly activated (Fig. S9G-I). These results suggested that DIM-5, DIM-7, and DIM-9 played similar roles in maintaining the state of the *lacO* array, consistent with them being members of one protein complex.

A more nuanced outcome was observed regarding the function of the HCHC complex at the *lacO* array. While the loss of HDA1 largely recapitulated the loss of DCDC (Fig. S9G-I), the *lacO* array still featured high levels of hH3 and H3K9me3 without CHAP (Fig. S9G-I). Surprisingly, in this strain, the *lacO* array was also found to undergo RIQ (Fig. S9G-I). Genome-wide, the *hda1Δ* condition was associated with a moderate depletion of hH3 over AT-rich DNA; however, this effect was much less dramatic compared to that observed upon deleting *hda1+* in Strain B (Fig. S2C; Fig. S9I). These results suggested that, while HDA1 acted as a critical regulator of heterochromatin at the *lacO* array, CHAP was involved in fine-tuning the balance between heterochromatin formation and RIQ activation.

### At the *lacO* array, RIQ can be decoupled from heterochromatin assembly

Our results showed that the *lacO* array could activate RIQ, assemble heterochromatin, and sustain high levels of genetic instability. It was interesting to know if any of those properties could be decoupled. To this end, we identified two clonal lineages of Strain C with spontaneous *lacO* deletions of 5247 and 8263 bps (Fig. S11A). The corresponding truncated arrays were named *d1* and *d2*, respectively (Fig. S11A). We found that *lacO(d1)* but not *lacO(d2)* could still induce RIQ (Fig. S11B). However, once DIM-5 was provided by transformation, both *lacO(d1)* and *lacO(d2)* became strongly heterochromatinized (Fig. S11C), suggesting that the ability of the *lacO* array to activate RIQ could be decoupled from its ability to assemble heterochromatin.

In principle, *lacO(d1)* could trigger RIQ because it contained some sequence motifs not present in *lacO(d2)*. Alternatively, a threshold amount of *lacO* DNA could be required. To differentiate between these possibilities, we generated three *dim-5Δ* strains with overlapping 1.4-1.6 kbp fragments of *lacO(d1)* (Fig. S11D,E). Neither fragment triggered RIQ; however, they all formed heterochromatin equally well once DIM-5 was provided by transformation (Fig. S11F,G). These results suggested that the ability to activate RIQ represented an emergent property of the *lacO* array, possibly controlled by a threshold-dependent mechanism.

## DISCUSSION

The great diversity and abundance of repetitive DNA necessitated the evolution of various processes to keep this fraction of the genome in check. While some of these processes, including the KZFP-TRIM28-SETDB1 pathway in animals [1] and the RITS-CLRC relay in fission yeast [47], employ strong sequence-specific cues to target dispersed repeats, others become activated by a large number of repeats organized as tandem arrays [48]. Such processes were suggested to involve transcription-based mechanisms [48–50], homologous DNA-DNA pairing [16,51], and non-B-DNA structures [52,53]. Tandem repeats were also found to associate with diverse non-histone proteins, including transcription factors [48,54,55].

We have found that in *N. crassa*, transcriptional and post-transcriptional silencing can be induced *de novo* by the aberrant activities of two unrelated DNA binding proteins (a synthetic TetR repressor and the endogenous GATA factor NIT-2) at the *tetO* array. While transcriptional silencing was mediated by canonical constitutive heterochromatin, post-transcriptional silencing resembled quelling but still occurred in the absence of RAD51 or RAD52, two critical recombination factors. Therefore, this process was named recombination-independent quelling (RIQ). In both situations involving TetR-GFP or NIT-2, the *tetO* array featured perturbed chromatin, as evidenced by decreased nucleosome occupancy and high MNase sensitivity. This state was ameliorated by SAD-6, which was also needed to initiate all silencing at the *tetO* array. Moreover, the properties of perturbed chromatin associated with RIQ of the *tetO* array were also relevant for RIQ of native AT-rich DNA. Thus, the state of perturbed chromatin emerged as a general signal for silencing (Fig. 5G). Importantly, while the loss of nucleosomes was linked to all *de novo* silencing in this study, the exact nature of the inducing signal remains unknown. Other candidate signals include R-loops and non-canonical DNA structures, which are all expected to correlate with nucleosome occupancy.

At the *tetO* array, heterochromatin and RIQ are initiated concomitantly yet independently from one another. In principle, with respect to heterochromatin assembly, the role of SAD-6 may be similar to the role of ATRX in the deposition of histone H3.3, which also requires DAXX (although no fungal ortholog of DAXX has been identified) [11]. On the other hand, the role of SAD-6 in RIQ appears profoundly mysterious.

### Recombination-independent quelling (RIQ) is a distinct RNAi process in *N. crassa*

Apart from quelling and MSUD, *N. crassa* sports two other RNAi pathways that can dynamically designate novel sRNA-producing loci [56]. The first (qiRNA) pathway is linked to DNA damage and requires RAD51 and RAD52, as well as the classical quelling factors QDE-1 and QDE-3 [56]. The second (disiRNA) pathway targets non-repetitive loci associated with sense/antisense transcription, does not require QDE-1 and QDE-3 [35], but depends on the exonuclease ERI1 [34]. Our results set these processes apart from RIQ (which needs QDE-1 and QDE-3 but not RAD51, RAD52, or ERI1). Yet RIQ and canonical quelling share the requirement for DNA replication and the H3K56 acetyltransferase RTT109 (which is also linked to DNA replication) [37]. In the case of RIQ, replication may be required to generate nascent chromatin [57], which may be particularly prone to perturbation by the aberrant activity of some DNA binding proteins (such as TetR-GFP and NIT-2).

Several types of RIQ appear to exist with respect to the requirement for SAD-6. While RIQ of the *tetO* array induced by TetR-GFP is absolutely dependent on SAD-6, low levels of *tetO*-derived sRNAs are still observed without SAD-6 upon nitrogen starvation. Further, RIQ of AT-rich DNA is only partially dependent on SAD-6, and RIQ of the *lacO* array occurs normally without SAD-6, implying that SAD-6 can be replaced by another factor (altogether, the *N. crassa* genome encodes 24 ATPases with predicted chromatin-remodeling functions [58]). Nevertheless, all types of RIQ involve perturbed chromatin, require QDE-1, and occur without RAD51, the only RecA-like recombinase in *N. crassa* [58]. All types of RIQ must also rely on transcription to produce single-stranded RNA that can be used as a template for synthesizing double-stranded RNA. While QDE-1 was proposed to mediate this step in the qiRNA pathway [59], the nature of the DNA-dependent RNA polymerase at the basis of RIQ remains unknown.

In *Neurospora*, two peculiar populations of sRNAs were previously described: one population corresponds to subtelomeric sRNAs, the other comprises sRNAs expressed at low background levels throughout the genome [35]. Notably, we have found that the levels of background sRNAs were apparently increased in all conditions linked to replication and recombination defects. In a subset of these conditions (specifically, without RAD51, RAD52, or RTT109), the levels of subtelomeric sRNAs became elevated as well. Neither of these effects was associated with the loss of QDE-3, consistent with the idea that QDE-3 plays a redundant role in DNA repair [60].

### Role of chromatin remodeling in the *de novo* initiation of silencing

The role of ATP-dependent remodelers in heterochromatin (re)assembly has been well established, with some archetypal examples including (in addition to ATRX-DAXX) the nucleolar remodeling complex (NoRC) [61], the nucleosome remodeling and deacetylase (NuRD) complex [62], Snf2/Hdac-containing repressor complex (SHREC) [63], as well as the helicases SMARCAD1 [64], HELLS/LSH [65] and its plant counterpart DDM1 [66,67]. These factors make nucleosomes more accessible or replaced altogether to promote the incorporation of histone variants or modifications favoring heterochromatin. Our results suggest that the aberrant activity of DNA binding proteins can produce a state of perturbed chromatin that may constitute a particularly favorable substrate for the remodeling-dependent silencing. Curiously, the occurrence of diverse transcription factors at mammalian pericentromeres was reported [48,54,55]. While some of these factors could induce the silencing directly, by recruiting additional enzymatic activities or driving transcription of non-coding RNAs [54], others may do so indirectly, by producing a state of perturbed chromatin. In budding yeast, the tight binding of LacI and TetR repressors to their operator arrays was shown to trigger transcriptional silencing [30,31]. According to our data, this process (i) can be induced by other DNA binding proteins, (ii) requires chromatin remodeling, and (iii) may also initiate post-transcriptional silencing.

How does SAD-6 recognize perturbed chromatin? Because SAD-6 controls RIQ of diverse loci (*i.e.*, the *tetO* array versus AT-rich DNA), it is unlikely to rely on sequence-specific signals or secondary structures that may recruit its orthologs in other situations [68]. On the other hand, SAD-6-dependent RIQ is associated with low nucleosome occupancy and high MNase sensitivity, suggesting that SAD-6 may favor nucleosome-free DNA. This propensity is shared by metazoan ATRX, which also localizes to nucleosome-depleted regions, including heavily transcribed genes [12], free proviral DNA [69], and R-loops [13]. In human cells, 87% of ATRX sites are present in open chromatin [70], and ATRX was also shown to interact with free nucleic acids *in vitro* [71]. Yet the exact nature of the SAD-6 recruitment mechanism and its relationship with the initiation of silencing all remain to be established.

In this study, the state of perturbed chromatin was induced by the aberrant activities of non-histone proteins at the *tetO* array. Yet, an analogous state might also be attained by other means, for example, by recombination-independent homologous pairing [72]. This idea is supported by the role of SAD-6 in MSUD, which relies on such pairing to find gaps in sequence identity between homologous chromosomes [20,22]. Thus, the proposed model (Fig. 5G), although derived from a synthetic experimental system, may also underpin other instances of the *de novo* initiation of transcriptional and post-transcriptional silencing in eukaryotes.

## ACKNOWLEDGMENTS

The work was supported by Agence Nationale de la Recherche (ANR-10-LABX-0062, ANR-11-LABX-0044, ANR-10-IDEX-0001-02, ANR-19-CE12-0002) and Institut Pasteur. We also acknowledge the Cell and Tissue Imaging Platform “PICT-IBiSA” (the Pasteur Imaging Facility, Institut Curie, part of the France Bioimaging National Infrastructure, funded by ANR-10-INBS-04).

## AUTHOR CONTRIBUTIONS

F.C. and E.G. designed research; F.C., S.C.R., J.K., S.C., I.L., and E.G. performed research; E.G. contributed new analytic tools; F.C., I.L., A.T., and E.G. analyzed data; E.G. wrote the paper.

## DATA, MATERIALS, AND SOFTWARE AVAILABILITY

All NGS data produced in this study were submitted to the Short Read Archive (BioProject PRJNA1109611). All strains created and analyzed in this study were submitted to the Fungal Genetics Stock Center (accessions 27332-27373). Custom genome references created and used in this study are available at figshare.com (DOI: 10.6084/m9.figshare.24943173.

## MATERIALS AND METHODS SUMMARY

A combination of ChIP-seq, MNase-seq, and sRNA-seq approaches was used to analyze the state of synthetic repetitive loci (represented by the *lacO* and the *tetO* operator arrays) and native AT-rich DNA. The following processes were assayed: (i) formation of constitutive heterochromatin, (ii) histone occupancy, (3) sensitivity to MNase, and (4) sRNAs expression.

## SUPPORTING INFORMATION (SI)

### MATERIALS AND METHODS

#### Plasmids

Plasmids were made by standard molecular cloning and validated by Sanger sequencing. Plasmid pFOC104A (carrying *lacO–tetO*) was made by inserting the NheI*–*XbaI fragment of pLAU43 [32] into pTSN6 [29]. This plasmid was propagated in *E. coli* DH10B cells in the presence of gentamicin and kanamycin [32]. All gene-deletion plasmids contained a marker conferring resistance to nourseothricin (clonNAT). Maps of all plasmids used in this study are provided in SI Data File 1.

#### Manipulation of *Neurospora* strains

##### A Strains

All *Neurospora* strains used in this study are listed in Table S2. All genes analyzed in this study are described in Table S1.

##### B Growth Media

Vogel’s minimal medium N with 1.5% sucrose (’VM medium’) was used for standard vegetative growth. The same medium but lacking ammonium nitrate (’VM-N medium) was used to induce acute nitrogen starvation. If needed, these media were supplemented with 100 μg/μl tetracycline, 0.1 μg/μl blasticidin S (to increase the expression of FLAG-SAD-6), or 0.1M hydroxyurea. Colonial growth was produced on sorbose agar (1x VM salts, 3% agar, 2% sorbose, 0.1% dextrose). For crossing, Synthetic Cross medium was used [73], containing 2% sucrose, 0.2 μg/ml biotin, and 2% agar. Crosses were setup on agar slants in glass culture tubes (20 x 200 mm). Ejected ascospores were collected in drops of water, induced by heat (60 °C for 30 min), and plated on sorbose agar at appropriate dilutions.

##### C Transformations

20 μl of macroconidia (washed and pelleted in 1M sorbitol) were combined with 1-2 μg of linearized plasmid DNA (dissolved in 10 μl of 1M sorbitol), incubated on ice for 20 min, and electroporated using the following settings: 1500 V, 600 Ω, 25 μF, 2 mm gap. Immediately following electroporation, macroconidia were mixed with 1 ml of 1M sorbitol. When transformed with plasmids targeting *his-3* or *csr-1*, macroconidia were plated on sorbose agar without delay. When transformed with plasmids carrying the nourseothricin resistance gene, macroconidia were allowed to recover for 4-5 hours at room temperature prior to being plated on sorbose agar supplemented with 50 μg/ml nourseothricin (Jena Bioscience, cat. no. AB-101L). Colonies were picked using glass Pasteur pipettes. Genomic DNA was prepared as previously described [74], and integration events were analyzed by PCR. Homokaryotic strains were obtained from primary transformants by macroconidiation. The integrity of the *lacO–tetO* locus was verified using the Expand Long Template PCR System (Roche) with the following primer pairs: P1/P3, P2/P5, P4/P7, and P6/P8 (Table S3). The *tetO* locus was verified analogously, using the primer pairs P1/P7 and P6/P8 (Table S3).

#### Assaying genetic stability of the arrays by multiplex PCR

Genomic DNA was extracted as previously described [74]. Two separate master mixes were prepared. Both mixes contained primers P9 and P10 (Table S3), which amplified a portion of the *spo11+* gene (as a positive control). In addition, the first master mix included primers P2 and P3 (to amplify the central part of the *lacO* array), while the second master mix included primers P6 and P7 (to amplify the central part of the *tetO* array). PCR products were resolved on 1.5% agarose gels and imaged using Gel Doc EZ (Bio-Rad).

#### Standard nitrogen starvation protocol

Each culture was started by inoculating thawed macroconidia (∼1 million CFUs) into 150 ml of VM medium. Cultures were grown for 24 hours at 30 °C/180 RPM. Resultant mycelia were transferred into chilled vacuum filtration units and rinsed 5 times with 100 ml of chilled water. Each mycelial sample was placed into 150 ml of VM-N medium and incubated for 5 hours at 30 °C/180 RPM. Mycelia were collected by filtration and split into two halves. One half was crosslinked using formaldehyde (described below), the other half was squeeze-dried and frozen in liquid nitrogen immediately.

#### Nitrogen starvation protocol including treatment with hydroxyurea (HU)

Each culture was started by inoculating thawed macroconidia (∼1 million CFUs) into 130 ml of VM medium. Cultures were pre-grown for 19 hours at 30 °C/180 RPM, adjusted to 130 g, supplemented with 20 ml of VM medium containing 1.14 g of HU, and returned to growth for additional 5 hours at 30 °C/180 RPM. Resultant mycelia were transferred into chilled vacuum filtration units, rinsed 5 times with 100 ml of chilled water, and returned to growth (5 hours, 30 °C/180 RPM) either in 150 ml of pure VM-N medium or in 150 ml of VM-N medium supplemented with 0.1M HU. Mycelia were collected by filtration and split into two halves. One half was crosslinked using formaldehyde (described below), the other half was squeeze-dried and frozen in liquid nitrogen immediately.

#### Chromatin immunoprecipitation (ChIP)

Each culture was started by inoculating thawed macroconidia (∼1 million CFUs) into 150 ml of VM medium. Cultures were grown for 24 hours at 30 °C/180 RPM. Resultant mycelia were collected by filtration, washed with phosphate-buffered saline (PBS) at room temperature, squeeze-dried, and transferred into 100 ml of PBS containing 1% formaldehyde. Crosslinking proceeded for 30 min at room temperature with constant stirring. After quenching with glycine for 5 min, mycelia were collected by filtration, rinsed with PBS, squeeze-dried, ground to fine powder in liquid nitrogen, and stored at −80 °C. The same mycelial prep was used for ChIP-seq and MNase-seq experiments.

For each sample, 150 mg of mycelial powder were combined with 1 ml of lysis buffer [140 mM NaCl; 1 mM EDTA; 1% Triton X-100; 0.1% sodium deoxycholate; 50 mM HEPES pH 7.5] containing 20 μl of proteinase inhibitors (Roche, cat. no. 11873580001, one tablet dissolved in 1 ml of PBS). Samples were transferred into 15-ml TPX tubes and sonicated in Bioruptor Plus (Diagenode) with the following settings: high power mode, 20 cycles, 90/90 sec ON/OFF, 4 °C. After sonication, samples were centrifuged twice for 5 min at 13000 ×g, 4 °C. SureBeads Protein A Magnetic Beads (Bio-Rad) were used for immunoprecipitation. Concentrated beads were washed and diluted five-fold in lysis buffer. The workflow included the following steps. First, each ChIP sample was pre-cleared using 20 μl of diluted beads (4 hours at 4 °C). Second, 25 μl of each pre-cleared ChIP sample were saved as ‘Input’. Third, an appropriate antibody (0.5-1.0 μl) was added and allowed to recognize its targets overnight (at 4 °C). The following antibodies were used: anti-FLAG (Sigma F1804), anti-H3K9me3 (Active Motif AB_2532132), anti-hH2B (Abcam ab1790), anti-hH3 (Abcam ab1791), and anti-GFP (Abcam ab290). Fourth, 20 μl of diluted beads, washed in lysis buffer containing 0.1% Tween-20, were added to each sample and incubated for 4 hours at 4 °C (all incubations were done using a rotating wheel). Fifth, beads were washed in the following buffers (1 ml, for 10 min): [i] lysis buffer (twice), [ii] lysis buffer with 0.5M NaCl, [iii] wash buffer [250 mM LiCl; 0.5% IGEPAL; 0.5% sodium deoxycholate; 1 mM EDTA; 10 mM TrisHCl pH 8.0], and {iv} TE buffer [1 mM EDTA; 10 mM TrisHCl pH 8.0]. Elution was done using TES buffer [1% SDS; 10 mM EDTA; 50 mM TrisHCl pH 8.0], for 10 min at 65 °C. Input fractions were also treated with TES buffer in parallel. Fractions were decrosslinked at 65 °C overnight, treated with RNAse A and then Proteinase K. DNA was purified by phenol-chloroform extraction, precipitated with ethanol, resuspended in 20 μl of TE buffer, and quantified using the Qubit dsDNA HS assay kit (Invitrogen). Two separate cultures (representing independent biological replicates) were analyzed for each condition.

#### MNase sensitivity analysis

For each sample, 80 mg of crosslinked mycelial powder were combined with 1 ml of lysis buffer including 20 μl of proteinase inhibitors (prepared as described above) and 2 mM CaCl_2_. Samples were mixed, equilibrated on ice for 10 min, mixed again, supplemented with 10 μl of MNase (NEB, cat. no. M0247S), and incubated in ThermoMixer C (Eppendorf) for 30 min at 37 °C. Reactions were stopped by adding 30 μl of 0.5M EGTA pH 8.0. Samples were centrifuged twice for 10 min at 16000 ×g, 4 °C. For decrosslinking, 50 μl of each sample were combined with 200 μl of TES buffer and incubated at 65 °C overnight, after which samples were treated with RNAse A and then Proteinase K. DNA was purified analogously to the ChIP protocol, and resuspended in 30 μl of TE buffer. DNA concentrations were measured using the Qubit dsDNA HS assay kit (Invitrogen).

#### Preparing ChIP-seq and MNase-seq libraries

ChIP-seq and MNase-seq libraries were made using the NEBNext Ultra II DNA Library Prep Kit for Illumina (NEB, cat. no. E7645S) and Agencourt Ampure XP beads (Beckman Coulter, cat. no. A63880), following the manufacturer’s instructions. Depending on the amount of starting material, libraries were amplified with 10-11 cycles of PCR. All ChIP-seq and MNase-seq libraries created and analyzed in this study are listed in Table S4 and Table S5, respectively.

#### Isolating small RNAs

Total RNA was extracted from ground mycelia with TRI Reagent (Sigma-Aldrich) and dissolved in 5M urea. RNA with a RINe score above 9.5 (as determined by the 4150 TapeStation System) was fractionated on 8% TBE-Urea gels. Approximately 60 μg of RNA were loaded on each gel, distributed in 6 wells. Populations of small RNAs in the 17-26 nt range were excised as gel slices. The latter were crushed and incubated overnight in 0.3 M NaCl at 25 °C/850 RPM. Gel debris was removed using 0.22-μm spin columns (Sigma-Aldrich, cat. no. CLS8161). Small RNAs were precipitated by adding 4 volumes of ethanol and centrifuging for 40 min at 20000 ×g, 4 °C. Pellets were dissolved in the total volume of 12 μl of water (per gel). Two separate cultures (representing independent biological replicates) were analyzed for each condition.

#### Preparing sRNA-seq libraries

sRNA-seq libraries were made as previously described [20]. For each library, 5 μl of sRNA sample were used, corresponding to 25 µg of input RNA (however, because a fraction of sRNAs was lost during purification, the amount of total RNA corresponding to each sRNA sample could be less). All sRNA-seq libraries created and analyzed in this study are listed in Table S6.

#### Sequencing libraries

All libraries were analyzed on the Agilent 4150 TapeStation System using D5000 and D1000 ScreenTapes for ChIP-seq/MNase-seq and sRNA-seq samples, respectively. 48 libraries were combined and sequenced on the NextSeq 500 machine (Illumina) using the High Output Kit v2.5 (75 cycles, single-end mode). Typically, 460-500 million indexed reads were generated, corresponding to the average library size of 10 million reads. Raw sequencing data were demultiplexed using bcl2fastq (v2.20) with ‘--barcode-mismatches 0 --no-lane-splitting’ flags. Adapter sequences were removed using cutadapt (v1.15) [75].

#### Designing custom genome references

Custom genome reference *mNC12* was created by concatenating the following GenBank entries: NC_026614.1, part of M54787.1 encoding the *mat a* locus (3237 bp), NC_026501.1, NC_026502.1, NC_026503.1, NC_026504.1, NC_026505.1, NC_026506.1, NC_026507.1, NW_011929459.1, NW_011929460.1, NW_011929461.1, NW_011929462.1, NW_011929463.1, NW_011929464.1, NW_011929465.1, NW_011929466.1, NW_011929467.1, NW_011929468.1, NW_011929469.1, NW_011929470.1, NW_011929471.1.

References *mNC12.lacO-tetO* and *mNC12.tetO* (SI Data File 2) were created based on *mNC12* by editing the sequence between *his-3* and *lpl* to reflect the structure of the locus after transformation with pFOC104A and pTSN6, respectively.

#### Sequence alignment and post-processing

All sequences were aligned to *mNC12.lacO-tetO* as the standard reference. Sequences originating from strains with the lone *tetO* array were also aligned to *mNC12.tetO*, while sequences derived from strains carrying parts of the *lacO* array were aligned to custom genome references that reflected the actual structure of the *his-3–lpl* locus in those strains. Alignment was done using bowtie2 (v2.3.4.1) with a standard set of parameters defined by the ‘--very-sensitive’ option [76]. ChIP-seq/MNase-seq reads were aligned to the genome directly, whereas sRNA-seq reads were first aligned to the *mito_ribo* reference (provided in SI Data File 2), the unaligned reads were retained and aligned to the genome reference. The alignments were post-processed using samtools (v1.7) [77].

#### Building and analyzing genomic profiles

For each library, a genome profile was constructed by counting the number of reads mapped over a series of non-overlapping 500-bp tiles (defined using the reference *mNC12.lacO-tetO*). Reads were counted using the command ‘samtools view -c’. Profiles were normalized by the effective number of reads as follows. For ChIP-seq/MNase-seq samples, the effective number of reads was equal to the total number of reads minus the reads mapped to the mitochondrial genome. For sRNA-seq samples, both mitochondrial and ribosomal reads were subtracted. Profiles of biological replicates were averaged to produce the final profile used for analysis. High-resolution profiles were created using exactly the same approach, except that the tile size was set to 1 bp. For all logarithmic operations to generate scatter plots, a pseudo-count of 1 was added to each each profile value. For computing and plotting physical maps of read-count ratios, only profile values in the denominator were augmented by 1. Matching ChIP-seq ‘Input’ profiles were used to normalize and plot MNase-seq profiles. All graphs were produced using ggplot2 [78]. Euler diagrams were created using the eulerr R package (v7.0.0, https://CRAN.R-project.org/package=eulerr).

#### Fluorescence microscopy analysis

Cultures were started by inoculating thawed macroconidia (∼10^5^ CFUs) into 5 ml of VM medium and grown for 6-7 hours at 30 °C/160 RPM. For the HU treatment: 38 mg of HU were added to a designated 5-ml culture (for the final concentration of 0.1M), and the culture was incubated for additional 24 hours at 30 °C/160 RPM. For all cultures: a 1-ml sample was taken and centrifuged at 10000 ×g, the pellets were collected and spread on glass slides. Images were acquired on an inverted wide-field microscope (Nikon TE2000) equipped with a 100×/1.4 NA immersion objective and a Prime BSI Express sCMOS camera (Teledyne Photometrics) with 6.5×6.5 μm pixel size. A SPECTRA X Light Engine lamp (Lumencor) set to power 50, for 100 ms, was used to illuminate the samples. For each area of interest, stacks of 32 images were obtained, with a z-step of 200 nm. Raw image data are available upon request.

### SI FIGURE LEGENDS

**Figure S1.**
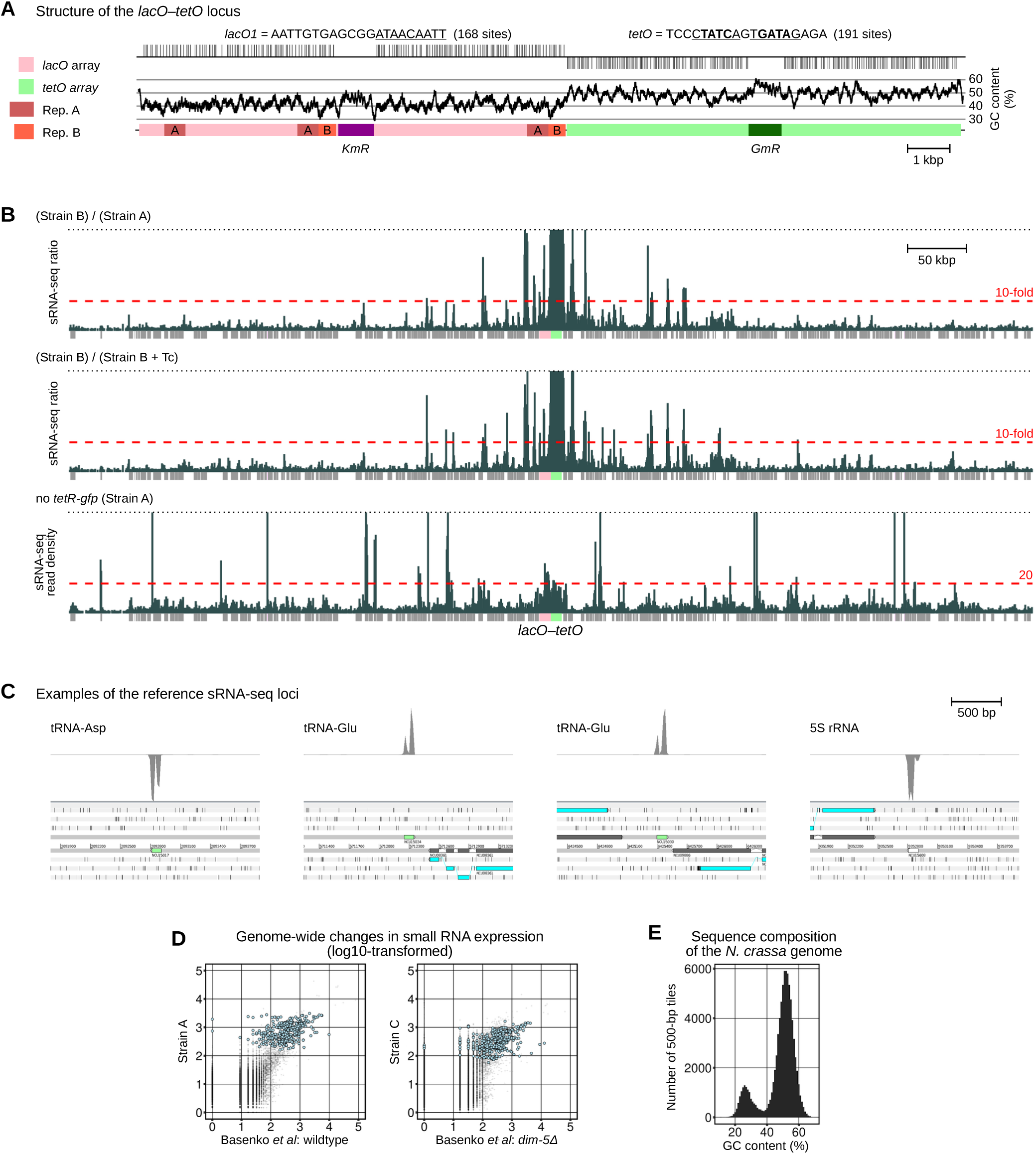
Experimental system. (A) Sequence properties of the *lacO–tetO* reporter are indicated (see text for explanation). Genes conferring resistance to kanamycin (*KmR*) and gentamicin (*GmR*) in *Escherichia coli* are also indicated. (B) Density of sRNA-seq reads in the 0.8-Mbp genomic region containing the *lacO–tetO* reporter (calculated per 500 bp, per 1 million mapped reads). Absolute and relative densities (corresponding to ratios of sRNA-seq counts) are shown, as indicated (plotted as in Fig. 5E). (C) Examples of the reference loci used to normalize and compare expression levels of array-derived sRNAs. For the purpose of the analysis, these loci were represented by 500-bp tiles (see text for explanation). (D) Scatter plots showing genome-wide changes in sRNA expression (analyzed and plotted as in Fig. 1F). Reference loci are shown as light-blue circles. ‘Basenko *et al*’ sRNA-seq data were published previously [40]. (E) Histogram showing composition of the *N. crassa* genome with respect to GC-content (analyzed using the same array of tiles as in Fig. 1F). AT-rich DNA is characterized by GC-content below 38-40%.

**Figure S2.**
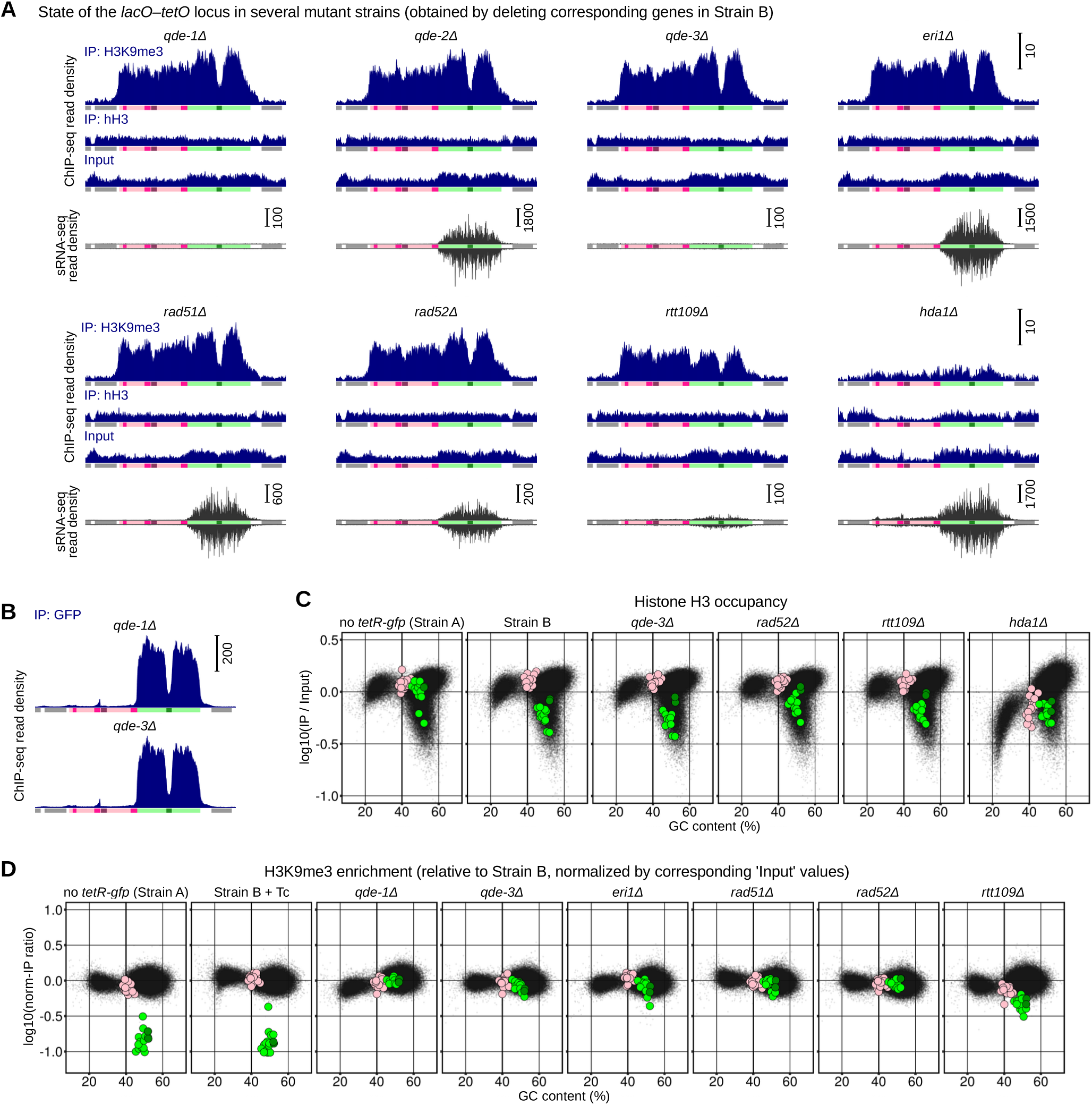
Silencing of the repressor-bound *tetO* array: properties and regulation. (A) Densities of ChIP-seq and sRNA-seq reads over the *lacO–tetO* reporter and the neighboring genes (analyzed and plotted as in Fig. 1D,E). (B) Density of ChIP-seq reads over the *lacO–tetO* reporter and the neighboring genes (analyzed and plotted as in Fig. 1D). (C) Scatter plots showing the relationship between the hH3 occupancy and GC-content (analyzed and plotted as in Fig. 2E). (D) Scatter plots showing the relationship between changes in H3K9me3 (normalized by their corresponding ‘Input’ values) and GC-content. Tiles overlapping the *lacO* and the *tetO* arrays are shown as pink and green circles, respectively.

**Figure S3.**
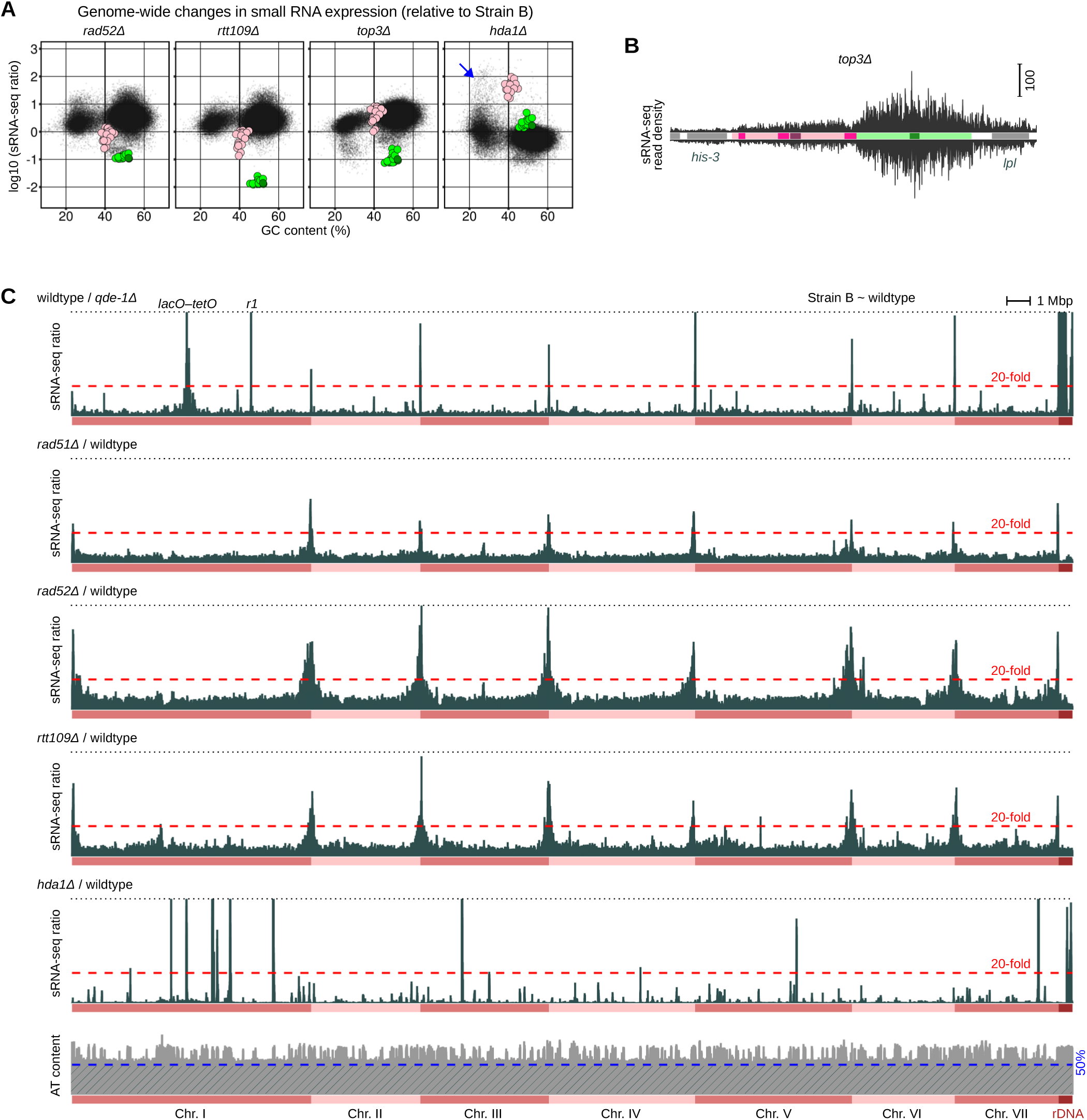
Genome-wide changes in sRNA expression. (A) Scatter plots showing the relationship between changes in sRNA expression and GC-content (analyzed and plotted as in Fig. 4C). Tiles overlapping the *lacO* and the *tetO* arrays are shown as pink and green circles, respectively. (B) Density of sRNA-seq reads over the *lacO–tetO* reporter and the neighboring genes (analyzed and plotted as in Fig. 1E). The increased expression of background sRNAs is evidenced by having multiple reads mapped to *his-3* and *lpl*. (C) Genome-wide changes in sRNA expression (analyzed and plotted as in Fig. 5E).

**Figure S4.**
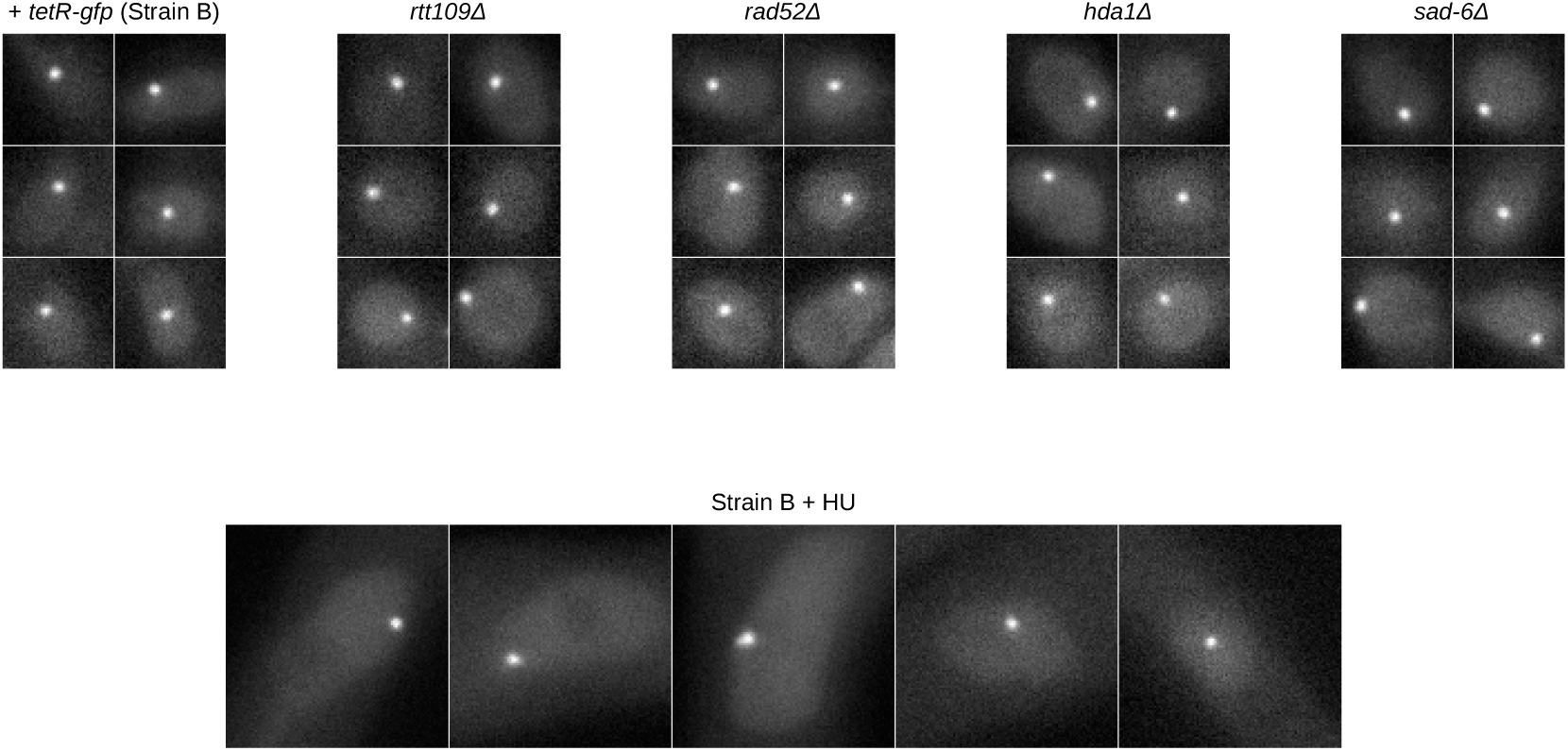
Fluorescence microscopy analysis of the repressor-bound *tetO* array. For each area of interest, image stacks were obtained with a z-step of 200 nm (Methods), and a single plane with the brightest localized signal was chosen for display. Images of several representative nuclei are shown for the indicated conditions.

**Figure S5.**
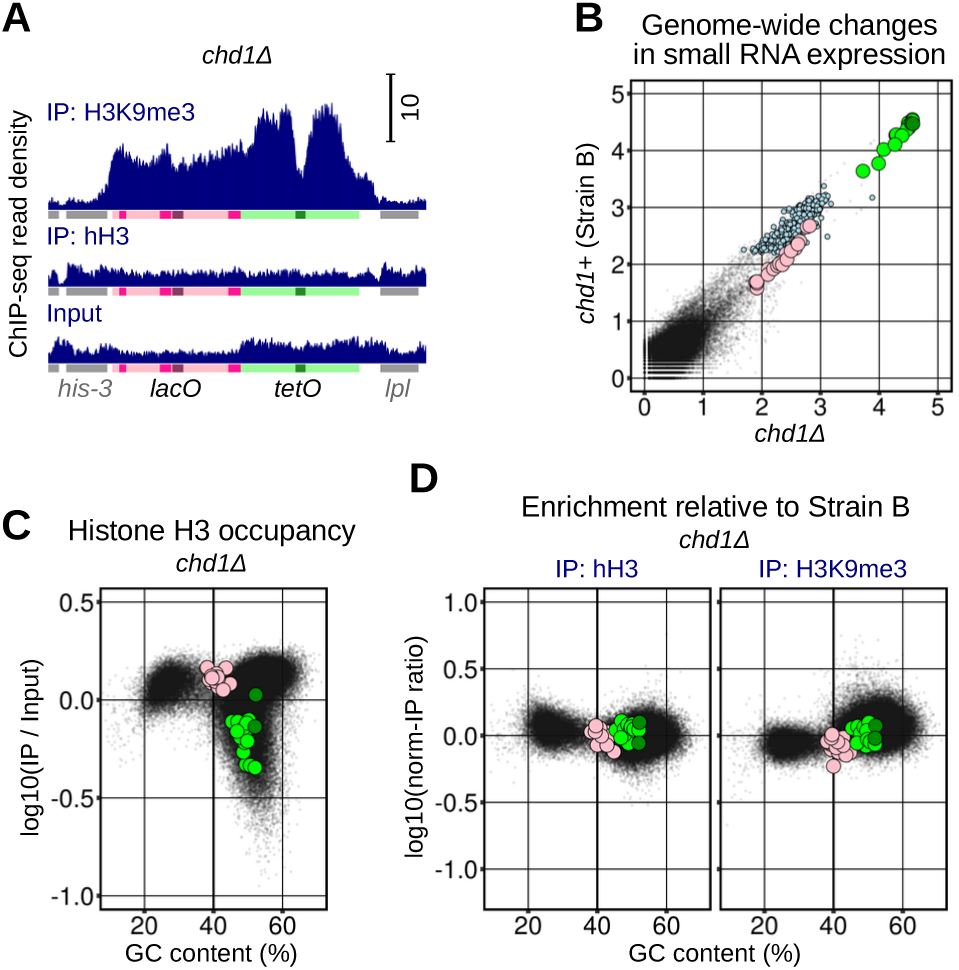
Silencing of the repressor-bound *tetO* array does not require CHD1. (A) Density of ChIP-seq reads over the *lacO–tetO* reporter and the neighboring genes (analyzed and plotted as in Fig. 1D). (B) Scatter plot showing genome-wide changes in sRNA expression (analyzed and plotted as in Fig. 1F). (C) Scatter plot showing the relationship between the hH3 occupancy and GC-content (analyzed and plotted as s in Fig. 2E). (D) Scatter plots showing the relationship between changes in hH3 occupancy and H3K9me3 versus GC-content (analyzed and plotted as in Fig. S2D).

**Figure S6.**
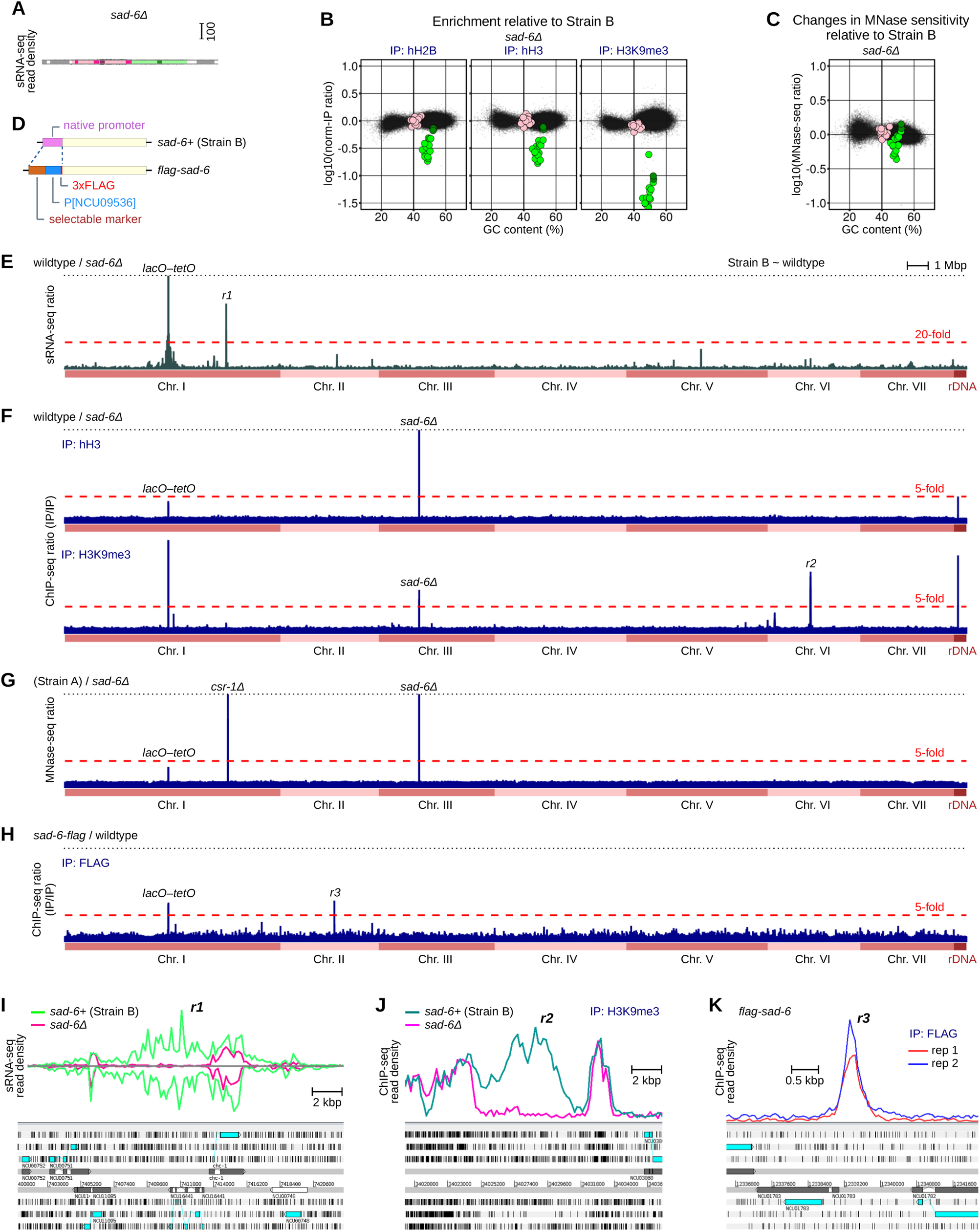
The role of SAD-6 in chromatin regulation. (A) Density of sRNA-seq reads over the *lacO–tetO* reporter and the neighboring genes (analyzed and plotted as in Fig. 1E). (B) Scatter plots showing the relationship between changes in histone occupancy and H3K9me3 versus GC-content (analyzed and plotted as in Fig. S2D). (C) Scatter plot showing the relationship between changes in MNase sensitivity and GC-content. Tiles overlapping the *lacO* and the *tetO* arrays are shown as pink and green circles, respectively. (D) Strategy to tag SAD-6 with 3xFLAG (the construct is provided on the plasmid pFOC109N). (E) Genome-wide changes in sRNA expression (analyzed and plotted as in Fig. 5E). (F) Genome-wide changes in hH3 occupancy and H3K9me3, calculated as ratios of ChIP-seq reads (the same array of 500-bp tiles was used as in Fig. 1F). Denominator values were augmented by 1. (G) Genome-wide changes in MNase sensitivity, calculated as ratios of MNase-seq reads (the same array of 500-bp tiles was used as in Fig. 1F). Denominator values were augmented by 1. (H) Genome-wide changes in FLAG-SAD-6 occupancy (analyzed and plotted as in Fig. S6F). (I) Region *r1*, in which sRNAs expression is strongly suppressed in the absence of SAD-6. (J) Region *r2*, in which constitutive heterochromatin is eliminated between two islands of AT-rich DNA in the absence of SAD-6. (K) Region *r3*, characterized by strong enrichment in FLAG-SAD-6.

**Figure S7.**
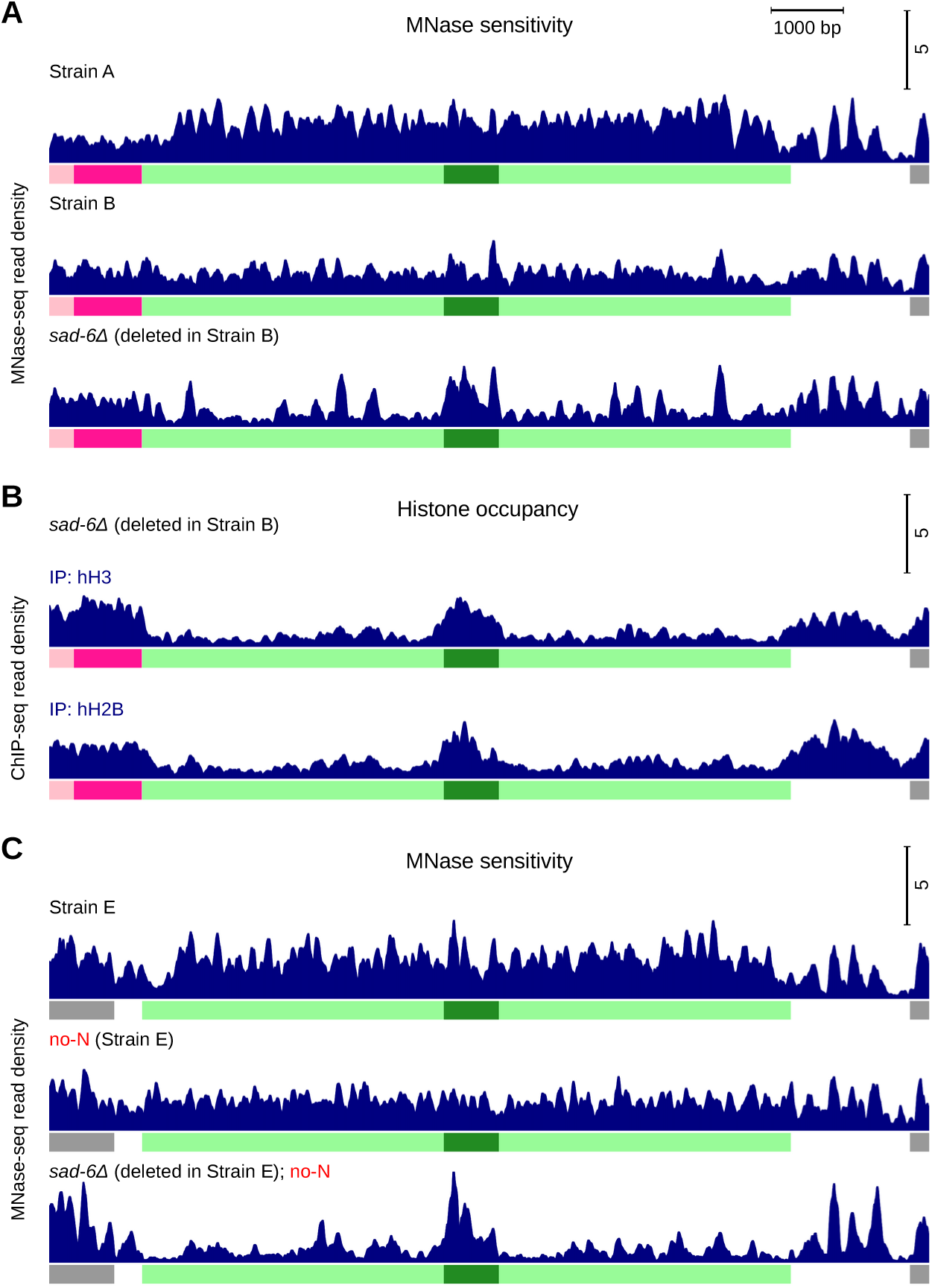
MNase-seq analysis of the *tetO* array. (A) Density of MNase-seq reads over the *tetO* array (analyzed and plotted as in Fig. 1D). (B) Density of ChIP-seq reads over the *tetO* array (analyzed and plotted as in Fig. 1D). (C) Density of MNase-seq reads over the *tetO* array (analyzed and plotted as in Fig. 1D).

**Figure S8.**
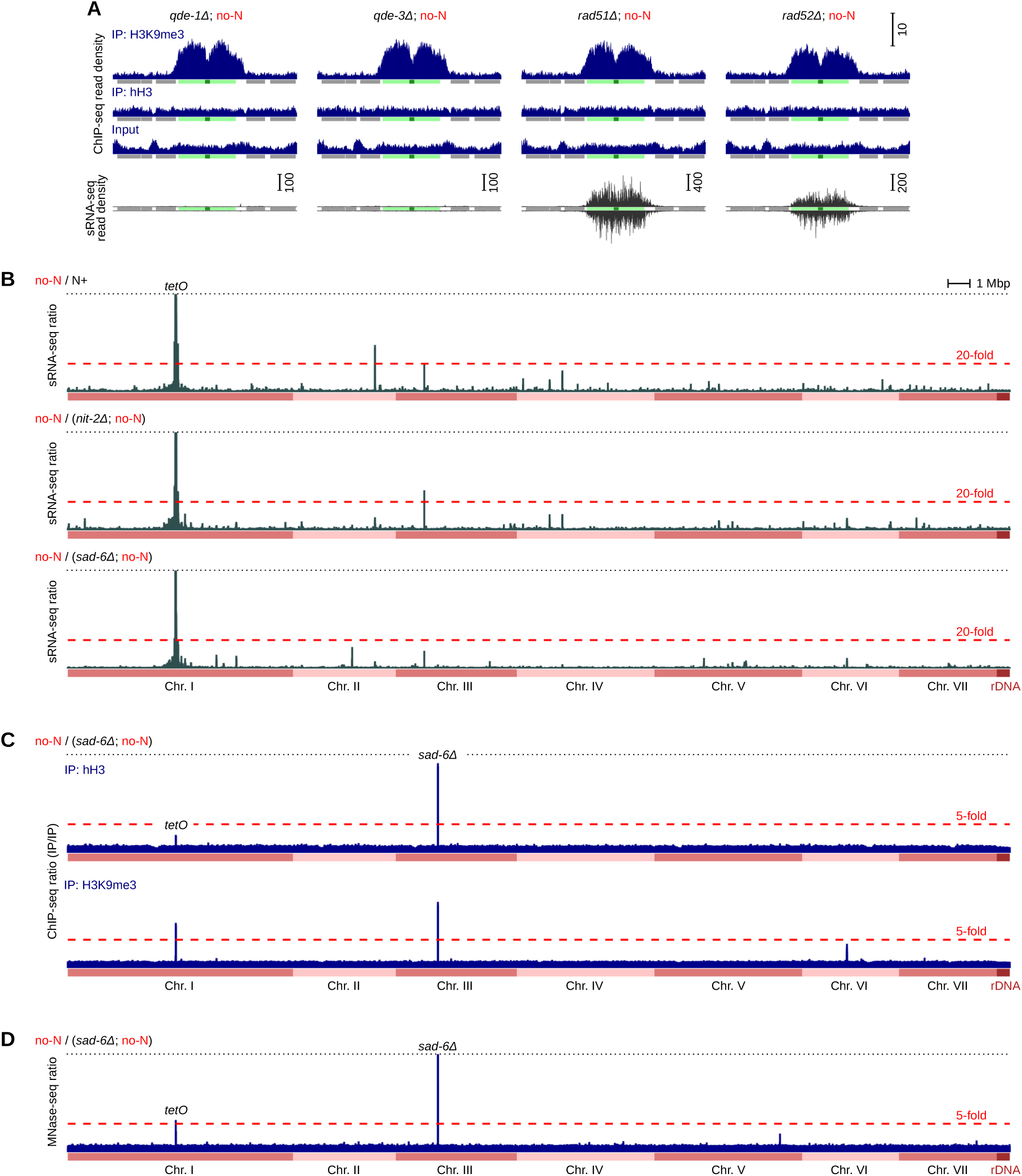
Starvation-induced silencing of the *tetO* array: properties and regulation. (A) Densities of ChIP-seq and sRNA-seq reads over the *tetO* array and the neighboring genes (analyzed and plotted as in Fig. 1D,E). (B) Genome-wide changes in sRNA expression (analyzed and plotted as in Fig. 5E). (C) Genome-wide changes in hH3 occupancy and H3K9me3 (analyzed and plotted as in Fig. S6F). (D) Genome-wide changes in MNase sensitivity (analyzed and plotted as in Fig. S6G).

**Figure S9.**
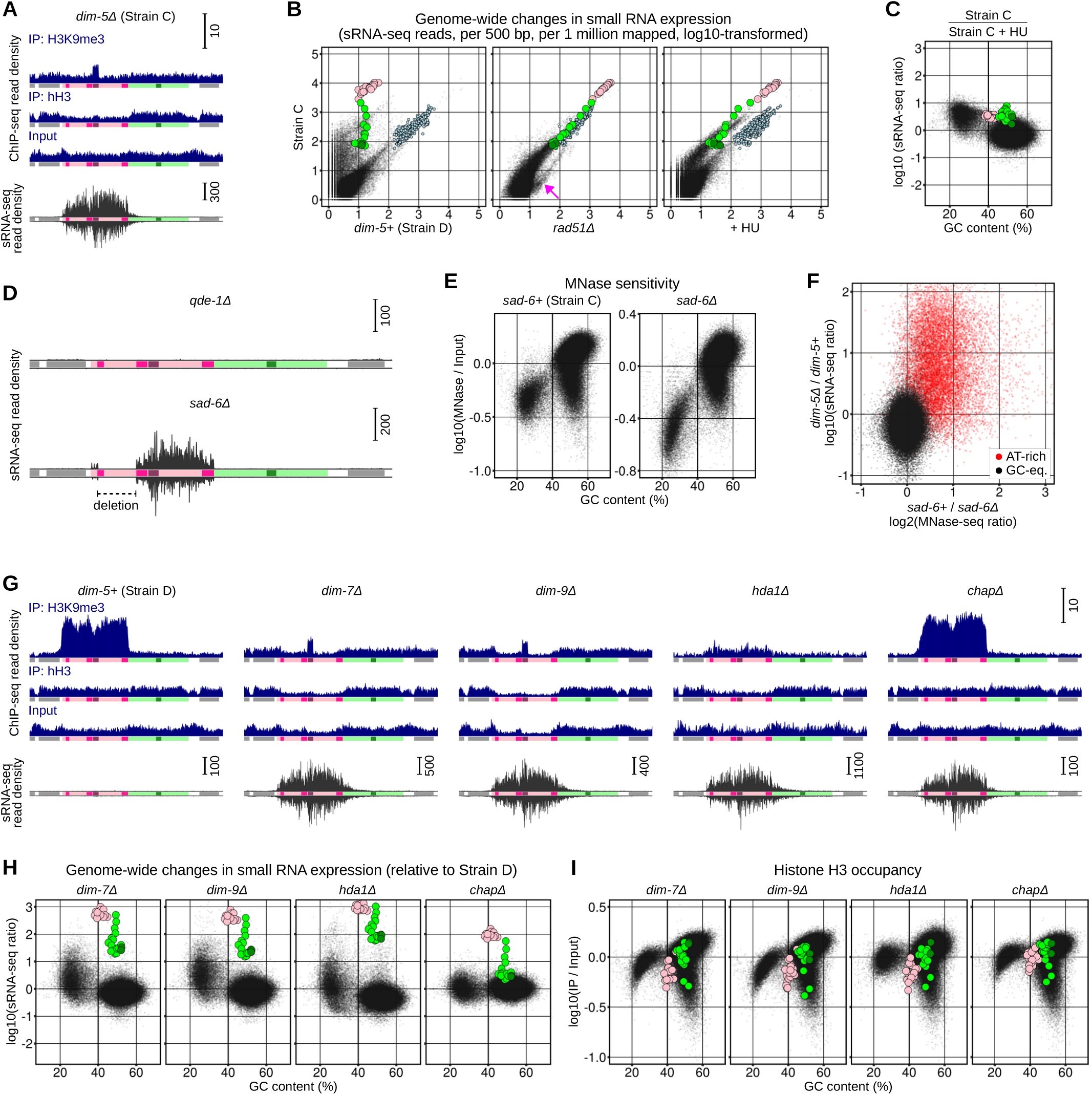
Post-transcriptional silencing of the *lacO* array and AT-rich DNA: properties and regulation. (A) Densities of ChIP-seq and sRNA-seq reads over the *lacO–tetO* reporter and the neighboring genes (analyzed and plotted as in Fig. 1D,E). (B) Scatter plots showing genome-wide changes in sRNA expression (analyzed and plotted as in Fig. 1F). Tiles overlapping the *lacO* and the *tetO* arrays are shown as pink and green circles, respectively. (C) Scatter plot showing the relationship between changes in sRNA expression and GC-content (analyzed and plotted as in Fig. 4C). Tiles overlapping the *lacO* and the *tetO* arrays are shown as pink and green circles, respectively. (D) Density of sRNA-seq reads over the *lacO–tetO* reporter and the neighboring genes (analyzed and plotted as in Fig. 1E). In the *sad-6Δ* strain (Strain ID: T752.13h, Table S2), the *lacO* array contained a large deletion. (E) Scatter plots showing the relationship between MNase sensitivity and GC-content (analyzed and plotted as in Fig. 2C). (F) Scatter plot showing the relationship between changes in sRNA expression (analyzed as in Fig. 4C) and changes in MNase sensitivity (analyzed as in Fig. 2C). AT-rich DNA is defined as in Fig. 5F). (G) Densities of ChIP-seq and sRNA-seq reads over the *lacO–tetO* reporter and the neighboring genes (analyzed and plotted as in Fig. 1D,E). (H) Scatter plots showing the relationship between changes in sRNA expression and GC-content (analyzed and plotted as Fig. 4C). Tiles overlapping the *lacO* and the *tetO* arrays are shown as pink and green circles, respectively. (I) Scatter plots showing the relationship between hH3 occupancy and GC-content (analyzed and plotted as in Fig. 2E).

**Figure S10.**
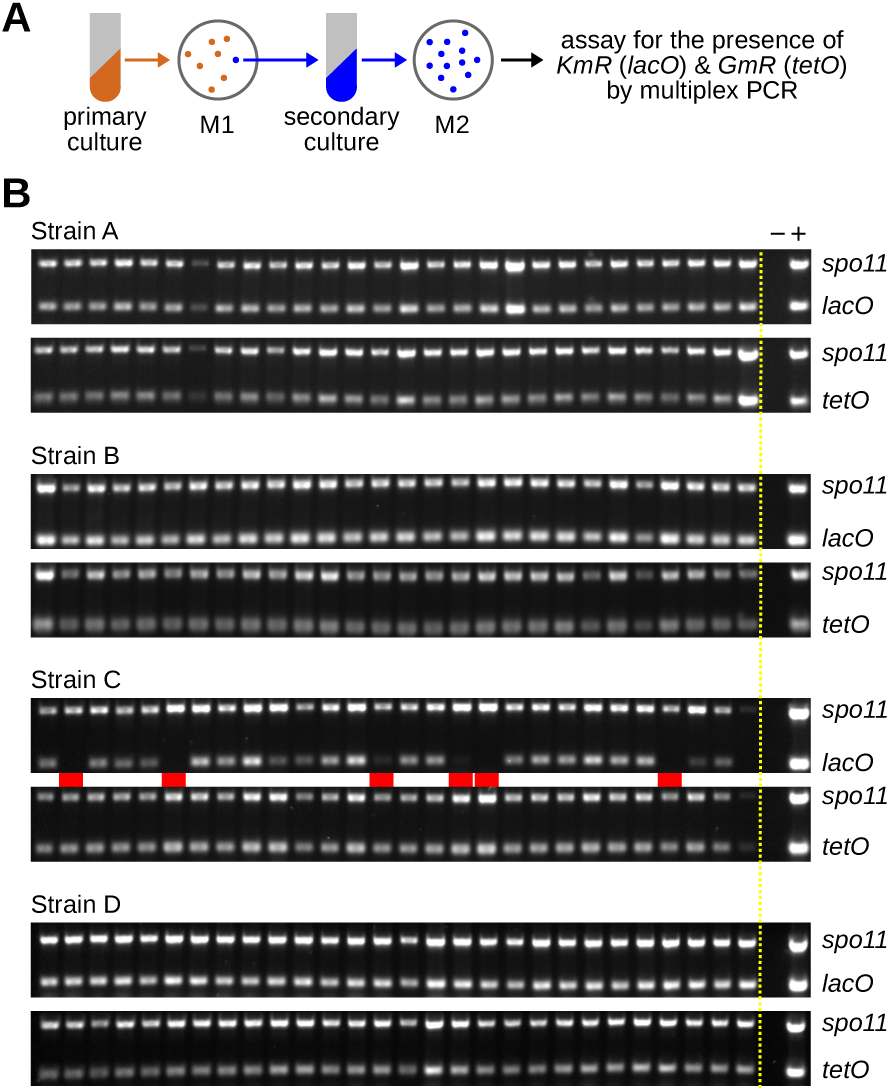
Assaying genetic stability of the *lacO–tetO* reporter. (A) Strategy to generate clonal populations of nuclei, in which the stability of the arrays was assayed. Several macroconidial isolates from a primary vegetative culture were tested by multiplex PCR for the presence of the arrays (see text for explanation). A single primary clone with the intact arrays was chosen to start a secondary culture, from which 28 random progeny were selected and re-assayed for the presence of the arrays. (B) PCR products were resolved on agarose gels. Missing products (corresponding to deletions) are indicated by red rectangles.

**Figure S11.**
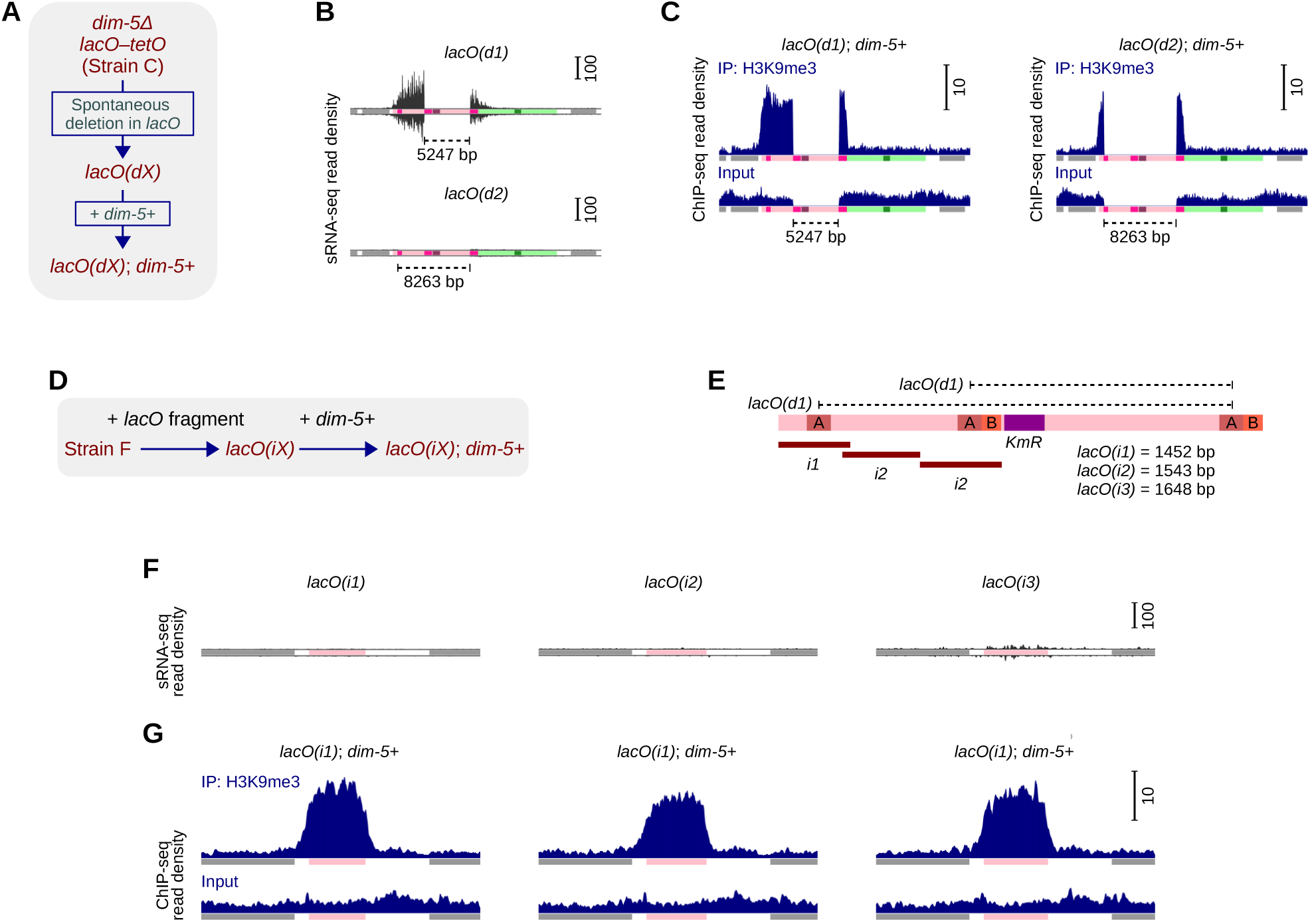
Transcriptional and post-transcriptional silencing of the *lacO* array can be decoupled. (A) A series of isogenic strains was produced with spontaneous deletions in the *lacO* array. In these strains, DIM-5 was either absent (original condition) or present (provided by transformation). (B) Density of sRNA-seq reads over the *lacO–tetO* reporter and the neighboring genes (analyzed and plotted as in Fig. 1E). Deletions are indicated. (C) Density of ChIP-seq reads over the *lacO–tetO* reporter and the neighboring genes (analyzed and plotted as in Fig. 1D). (D) A series of isogenic strains was produced with fragments of the *lacO* array. In these strains, DIM-5 was either absent (original condition) or present (provided by transformation). (E) Deletions and insertions of the *lacO* array analyzed in this study. (F) Density of sRNA-seq reads over the *lacO* fragment and the neighboring genes (analyzed as in Fig. 1E). (G) Density of ChIP-seq reads over the *lacO* fragment and the neighboring genes (analyzed as in Fig. 1D).

### SI TABLES

**Table S1.** *Neurospora* genes analyzed in this study.

| Gene | Standard name | Linkage group | Protein accession | Full name | Synonym (if any) |
| --- | --- | --- | --- | --- | --- |
| *chap* | NCU01796 | II | XP_956474.1 | *CDP-2- and HDA1 associating protein* |  |
| *chd1* | NCU03060 | I | XP_011393066.1 | *chromatin remodeling factor 6-1* | *crf6-1* |
| *csr-1* | NCU00726 | I | XP_011392822.1 | *cyclosporin-resistant-1* | *cyp* |
| *dim-2* | NCU02247 | VII | XP_959891.1 | *defective in methylation-2* |  |
| *dim-5* | NCU04402 | IV | XP_957479.2 | *defective in methylation-5* | *Su(var)3-9* |
| *dim-7* | NCU04152 | V | XP_961308.2 | *defective in methylation-7* |  |
| *dim-9* | NCU01656 | II | XP_956278.3 | *defective in methylation-9* |  |
| *eri1* | NCU06684 | V | XP_011394691.1 | *RNA binding protein* |  |
| *hda1* | NCU01525 | II | XP_956974.3 | *histone deacetylase-1* |  |
| *hh2b* | NCU02435 | VII | XP_959440.1 | *histone H2B* |  |
| *hh3* | NCU01635 | II | XP_956003.1 | *histone H3* |  |
| *his-3* | NCU03139 | I | XP_964188.1 | *histidine-3* |  |
| *lpl* | NCU03141 | I | XP_964190.2 | *lysophospholipase* |  |
| *mus-52* | NCU00077 | III | XP_956387.3 | *mutagen sensitive-52* | *KU80* |
| *nit-2* | NCU09068 | I | XP_963796.3 | *nitrate nonutilizer-2* | *areA* |
| *qde-1* | NCU07534 | III | XP_959047.1 | *quelling-defective-1* |  |
| *qde-2* | NCU04730 | VI | XP_011394903.1 | *quelling-defective-2* |  |
| *qde-3* | NCU08598 | I | XP_964030.3 | *quelling-defective-3* | *mus-19* |
| *rad51* | NCU02741 | I | XP_965126.1 | *meiotic-3* | *mei-3* |
| *rad52* | NCU04275 | V | XP_961065.2 | *mutagen sensitive-11* | *mus-11* |
| *rid* | NCU02034 | I | XP_011392925.1 | *RIP defective* |  |
| *rtt109* | NCU09825 | V | XP_958524.1 | *DNA damage response protein Rtt109* |  |
| *sad-6* | NCU06190 | III | XP_963002.3 | *atrx ortholog* | *ATRX* |
| *set-7* | NCU07496 | I | XP_965043.2 | *SET-domain histone methyltransferase-7* | *E(z)* |
| *spo11* | NCU01120 | V | XP_961027.3 | *spo11 ortholog* |  |
| *top3* | NCU00081 | III | XP_956391.3 | *DNA topoisomerase 3-beta* |  |

**Table S2.** *Neurospora* strains used in this study.

| Genetic condition | Strain ID | Genotype | Source |
| --- | --- | --- | --- |
| Strain O | C58.7 | *a; ridΔ; dim-2Δ; mus-52Δ; his-3* | ref. 16 |
| Strain A | T647.1h | *a; ridΔ; dim-2Δ; mus-52Δ; his-3+::lacO-tetO* | C58.7 transformed with pFOC104A |
| Strain B | T658.1 | *a; ridΔ; dim-2Δ; mus-52Δ; his-3+::lacO-tetO; csr-1::tetR-gfp* | T647.1h transformed with pTSN7B |
| Strain B::chd1Δ | T709.3h | *a; ridΔ; dim-2Δ; mus-52Δ; his-3+::lacO-tetO; csr-1::tetR-gfp; chd1Δ* | T658.1 transformed with pFOC111B |
| Strain B::eri1Δ | T770.2h | *a; ridΔ; dim-2Δ; mus-52Δ; his-3+::lacO-tetO; csr-1::tetR-gfp; eri1Δ* | T658.1 transformed with pSCR8B |
| Strain B::hda1Δ | T714.8h | *a; ridΔ; dim-2Δ; mus-52Δ; his-3+::lacO-tetO; csr-1::tetR-gfp; hda1Δ* | T658.1 transformed with pFOC121B |
| Strain B::qde-1Δ | T710.2h | *a; ridΔ; dim-2Δ; mus-52Δ; his-3+::lacO-tetO; csr-1::tetR-gfp; qde-1Δ* | T658.1 transformed with pFOC120B |
| Strain B::qde-2Δ | T711.21h | *a; ridΔ; dim-2Δ; mus-52Δ; his-3+::lacO-tetO; csr-1::tetR-gfp; qde-2Δ* | T658.1 transformed with pFOC113B |
| Strain B::qde-3Δ | T712.2h | *a; ridΔ; dim-2Δ; mus-52Δ; his-3+::lacO-tetO; csr-1::tetR-gfp; qde-3Δ* | T658.1 transformed with pFOC114B |
| Strain B::rad51Δ | T737.1h | *a; ridΔ; dim-2Δ; mus-52Δ; his-3+::lacO-tetO; csr-1::tetR-gfp; rad51Δ* | T658.1 transformed with pFOC106B |
| Strain B::rad52Δ | T738.1h | *a; ridΔ; dim-2Δ; mus-52Δ; his-3+::lacO-tetO; csr-1::tetR-gfp; rad52Δ* | T658.1 transformed with pFOC107B |
| Strain B::rtt109Δ | T786.3h | *a; ridΔ; dim-2Δ; mus-52Δ; his-3+::lacO-tetO; csr-1::tetR-gfp; rtt109Δ* | T658.1 transformed with pFOC129B |
| Strain B::sad-6Δ | T707.51h | *a; ridΔ; dim-2Δ; mus-52Δ; his-3+::lacO-tetO; csr-1::tetR-gfp; sad-6Δ* | T658.1 transformed with pFOC109B |
| Strain B::flag-sad-6 | T827.7h | *a; ridΔ; dim-2Δ; mus-52Δ; his-3+::lacO-tetO; csr-1::tetR-gfp; flag-sad-6* | T658.1 transformed with pFOC109N |
| Strain B::top3Δ | T812.11h | *a; ridΔ; dim-2Δ; mus-52Δ; his-3+::lacO-tetO; csr-1::tetR-gfp; top3Δ* | T658.1 transformed with pSCR12B |
| Strain E | T601.2h | *a; ridΔ; dim-2Δ; mus-52Δ; his-3+::tetO* | C58.7 transformed with pTSN6 |
| Strain E::tetR-gfp | T753.1 | *a; ridΔ; dim-2Δ; mus-52Δ; his-3+::tetO; csr-1::tetR-gfp* | T601.2h transformed with pTSN7B |
| Strain E::nit-2Δ | T851.1h | *a; ridΔ; dim-2Δ; mus-52Δ; his-3+::tetO; nit-2Δ* | T601.2h transformed with pEAG276B |
| Strain E::qde-1Δ | T839.10h | *a; ridΔ; dim-2Δ; mus-52Δ; his-3+::tetO; qde-1Δ* | T601.2h transformed with pFOC120B |
| Strain E::qde-3Δ | T840.10h | *a; ridΔ; dim-2Δ; mus-52Δ; his-3+::tetO; qde-3Δ* | T601.2h transformed with pFOC114B |
| Strain E::rad51Δ | T847.3h | *a; ridΔ; dim-2Δ; mus-52Δ; his-3+::tetO; rad51Δ* | T601.2h transformed with pFOC106B |
| Strain E::rad52Δ | T848.1h | *a; ridΔ; dim-2Δ; mus-52Δ; his-3+::tetO; rad52Δ* | T601.2h transformed with pFOC107B |
| Strain E::sad-6Δ | T774.4h | *a; ridΔ; dim-2Δ; mus-52Δ; his-3+::tetO; sad-6Δ* | T601.2h transformed with pFOC109B |
| Strain G | C252.1 | *A; ridΔ; dim-2Δ; dim-5Δ; set-7Δ; sad-2Δ* | Cross progeny of C205.11 & C244.9 |
| Strain C | C290.6 | *A; ridΔ; dim-2Δ; dim-5Δ; set-7Δ; mus-52Δ; his-3+::lacO-tetO* | Cross progeny of T647.1h & C252.1 |
| Strain C::lacO(d1) | C290.6.C4 | *A; ridΔ; dim-2Δ; dim-5Δ; set-7Δ; mus-52Δ; his-3+::lacO(d1)-tetO* | Clonal isolate of C290.6 |
| Strain C::lacO(d2) | C290.6.C6 | *A; ridΔ; dim-2Δ; dim-5Δ; set-7Δ; mus-52Δ; his-3+::lacO(d2)-tetO* | Clonal isolate of C290.6 |
| Strain C::lacO(d1); dim-5+ | T697.1 | *A; ridΔ; dim-2Δ; dim-5Δ; set-7Δ; mus-52Δ; his-3+::lacO(d1)-tetO; csr-1::dim-5+* | C290.6.C4 transformed with pEAG247A |
| Strain C::lacO(d2); dim-5+ | T698.1 | *A; ridΔ; dim-2Δ; dim-5Δ; set-7Δ; mus-52Δ; his-3+::lacO(d2)-tetO; csr-1::dim-5+* | C290.6.C6 transformed with pEAG247A |
| Strain C::dim-5+ | T663.2 | *A; ridΔ; dim-2Δ; dim-5Δ; set-7Δ; mus-52Δ; his-3+::lacO-tetO; csr-1::dim-5+* | C290.6 transformed with pEAG247A (Strain D) |
| Strain C::qde-1Δ | T739.4h | *A; ridΔ; dim-2Δ; dim-5Δ; set-7Δ; mus-52Δ; his-3+::lacO-tetO; qde-1Δ* | C290.6 transformed with pFOC120B |
| Strain C::rad51Δ | T678.5h | *A; ridΔ; dim-2Δ; dim-5Δ; set-7Δ; mus-52Δ; his-3+::lacO-tetO; rad51Δ* | C290.6 transformed with pFOC106B |
| Strain C::sad-6Δ | T752.13h | *A; ridΔ; dim-2Δ; dim-5Δ; set-7Δ; mus-52Δ; his-3+::lacO-tetO; sad-6Δ* | C290.6 transformed with pFOC109B |
| Strain D::chapΔ | T690.1h | *A; ridΔ; dim-2Δ; dim-5Δ; set-7Δ; mus-52Δ; his-3+::lacO-tetO; csr-1::dim-5+; chapΔ* | T663.2 transformed with pFOC118B |
| Strain D::dim-7Δ | T688.2h | *A; ridΔ; dim-2Δ; dim-5Δ; set-7Δ; mus-52Δ; his-3+::lacO-tetO; csr-1::dim-5+; dim-7Δ* | T663.2 transformed with pFOC116B |
| Strain D::dim-9Δ | T689.2h | *A; ridΔ; dim-2Δ; dim-5Δ; set-7Δ; mus-52Δ; his-3+::lacO-tetO; csr-1::dim-5+; dim-9Δ* | T663.2 transformed with pFOC117B |
| Strain D::hda1Δ | T743.3h | *A; ridΔ; dim-2Δ; dim-5Δ; set-7Δ; mus-52Δ; his-3+::lacO-tetO; csr-1::dim-5+; hda1Δ* | T663.2 transformed with pFOC121B |
| Strain F | C146.1 | *a; ridΔ; dim-5Δ; set-7Δ; mus-52Δ; his-3* | ref. 16 |
| Strain F::lacO(i1) | T724.2h | *a; ridΔ; dim-5Δ; set-7Δ; mus-52Δ; his-3+::lacO(i1)* | C146.1 transformed with pFOC125A |
| Strain F::lacO(i2) | T726.1h | *a; ridΔ; dim-5Δ; set-7Δ; mus-52Δ; his-3+::lacO(i2)* | C146.1 transformed with pFOC125C |
| Strain F::lacO(i3) | T728.1h | *a; ridΔ; dim-5Δ; set-7Δ; mus-52Δ; his-3+::lacO(i3)* | C146.1 transformed with pFOC125E |
| Strain F::lacO(i1); dim-5+ | T764.1 | *a; ridΔ; dim-5Δ; set-7Δ; mus-52Δ; his-3+::lacO(i1); csr-1::dim-5+* | T724.2h transformed with pEAG247A |
| Strain F::lacO(i2); dim-5+ | T766.1 | *a; ridΔ; dim-5Δ; set-7Δ; mus-52Δ; his-3+::lacO(i2); csr-1::dim-5+* | T726.1h transformed with pEAG247A |
| Strain F::lacO(i3); dim-5+ | T776.1 | *a; ridΔ; dim-5Δ; set-7Δ; mus-52Δ; his-3+::lacO(i3); csr-1::dim-5+* | T728.1h transformed with pEAG247A |

**Table S3.**
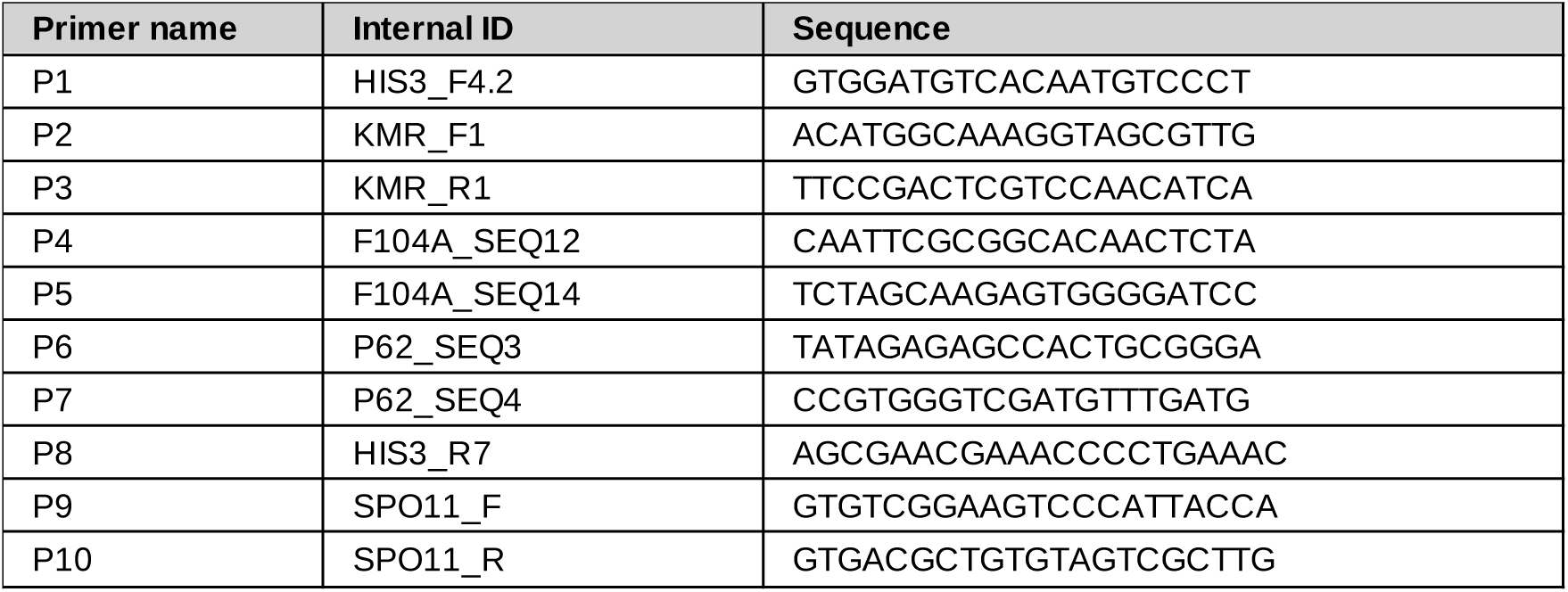
Diagnostic primers used in this study.

**Table S4.** ChIP-seq libraries created and analyzed in this study.

| # | Genetic condition | Strain ID | Library | IP (or Input) | Alignment rate (%) | Aligned reads * | Input library | Additional conditions |
| --- | --- | --- | --- | --- | --- | --- | --- | --- |
| 1 | Strain A | T647.1h | CD397_i17_A | GFP | 98.3 | 7929410 | CD394_i14_A |  |
| 2 | Strain A | T647.1h | CD593_i44_A | GFP | 87.2 | 8976252 | CD590_i48_A |  |
| 3 | Strain A | T647.1h | CD2_i3_A | H3K9me3 | 96.5 | 5257486 | CD1_i1_A |  |
| 4 | Strain A | T647.1h | CD396_i16_A | H3K9me3 | 97.4 | 7609395 | CD394_i14_A |  |
| 5 | Strain A | T647.1h | CD8_i5_A | hH3 | 99.0 | 7766288 | CD1_i1_A |  |
| 6 | Strain A | T647.1h | CD395_i15_A | hH3 | 98.9 | 7798269 | CD394_i14_A |  |
| 7 | Strain A | T647.1h | CD1_i1_A | Input | 99.0 | 5636932 | N/A |  |
| 8 | Strain A | T647.1h | CD394_i14_A | Input | 99.0 | 8815831 | N/A |  |
| 9 | Strain A | T647.1h | CD590_i48_A | Input | 99.0 | 11094748 | N/A |  |
| 10 | Strain B | T658.1 | CD669_i48_A | FLAG | 96.2 | 8815307 | CD529_i11_A |  |
| 11 | Strain B | T658.1 | CD671_i47_A | FLAG | 96.7 | 10365761 | CD594_i27_A |  |
| 12 | Strain B | T658.1 | CD463_i8_A | GFP | 97.3 | 8909320 | CD459_i43_A | Tc |
| 13 | Strain B | T658.1 | CD477_i10_A | GFP | 98.7 | 6396447 | CD473_i5_A | Tc |
| 14 | Strain B | T658.1 | CD28_i19_A | GFP | 96.9 | 10006244 | CD26_i17_A |  |
| 15 | Strain B | T658.1 | CD221_i6_A | GFP | 93.1 | 4276945 | CD220_i5_A |  |
| 16 | Strain B | T658.1 | CD461_i45_A | H3K9me3 | 97.3 | 8697139 | CD459_i43_A | Tc |
| 17 | Strain B | T658.1 | CD475_i7_A | H3K9me3 | 97.2 | 6597607 | CD473_i5_A | Tc |
| 18 | Strain B | T658.1 | CD29_i20_A | H3K9me3 | 97.6 | 10922524 | CD26_i17_A |  |
| 19 | Strain B | T658.1 | CD92_i43_A | H3K9me3 | 96.9 | 7653828 | CD89_i40_A |  |
| 20 | Strain B | T658.1 | CD731_i26_A | hH2B | 99.0 | 9636938 | CD459_i43_A | Tc |
| 21 | Strain B | T658.1 | CD732_i27_A | hH2B | 99.0 | 9454969 | CD473_i5_A | Tc |
| 22 | Strain B | T658.1 | CD729_i24_A | hH2B | 98.8 | 9106248 | CD529_i11_A |  |
| 23 | Strain B | T658.1 | CD730_i25_A | hH2B | 99.0 | 9851625 | CD594_i27_A |  |
| 24 | Strain B | T658.1 | CD460_i44_A | hH3 | 98.8 | 11649989 | CD459_i43_A | Tc |
| 25 | Strain B | T658.1 | CD474_i6_A | hH3 | 99.0 | 6252061 | CD473_i5_A | Tc |
| 26 | Strain B | T658.1 | CD27_i18_A | hH3 | 99.0 | 7549064 | CD26_i17_A |  |
| 27 | Strain B | T658.1 | CD90_i41_A | hH3 | 98.8 | 7775513 | CD89_i40_A |  |
| 28 | Strain B | T658.1 | CD459_i43_A | Input | 99.1 | 15466837 | N/A | Tc |
| 29 | Strain B | T658.1 | CD473_i5_A | Input | 99.0 | 10379300 | N/A | Tc |
| 30 | Strain B | T658.1 | CD26_i17_A | Input | 99.0 | 6046340 | N/A |  |
| 31 | Strain B | T658.1 | CD89_i40_A | Input | 98.9 | 8480540 | N/A |  |
| 32 | Strain B | T658.1 | CD220_i5_A | Input | 98.7 | 7247386 | N/A |  |
| 33 | Strain B | T658.1 | CD529_i11_A | Input | 98.9 | 8477778 | N/A |  |
| 34 | Strain B | T658.1 | CD594_i27_A | Input | 98.8 | 9366140 | N/A |  |
| 35 | Strain B::chd1Δ | T709.3h | CD182_i20_A | H3K9me3 | 97.3 | 10597328 | CD180_i18_A |  |
| 36 | Strain B::chd1Δ | T709.3h | CD441_i25_A | H3K9me3 | 97.7 | 10717376 | CD439_i23_A |  |
| 37 | Strain B::chd1Δ | T709.3h | CD181_i19_A | hH3 | 81.4 | 8090865 | CD180_i18_A |  |
| 38 | Strain B::chd1Δ | T709.3h | CD440_i24_A | hH3 | 99.0 | 9477493 | CD439_i23_A |  |
| 39 | Strain B::chd1Δ | T709.3h | CD180_i18_A | Input | 98.6 | 8172534 | N/A |  |
| 40 | Strain B::chd1Δ | T709.3h | CD439_i23_A | Input | 99.2 | 8324205 | N/A |  |
| 41 | Strain B::eri1Δ | T770.2h | CD240_i22_A | H3K9me3 | 96.5 | 8209183 | CD238_i20_A |  |
| 42 | Strain B::eri1Δ | T770.2h | CD431_i15_A | H3K9me3 | 97.5 | 9112162 | CD429_i13_A |  |
| 43 | Strain B::eri1Δ | T770.2h | CD430_i14_A | hH3 | 99.1 | 7200067 | CD429_i13_A |  |
| 44 | Strain B::eri1Δ | T770.2h | CD239_i21_A | hH3 | 98.3 | 7230932 | CD238_i20_A |  |

\* Number of aligned reads is reported for the nuclear genome (corresponding to the interval 64841-41123470 in *mNC12.lacO-tetO*)

**Table S4. ChIP-seq libraries created and analyzed in this study (continued).**
|  |  |  |  |  |  |  |  |  |
| --- | --- | --- | --- | --- | --- | --- | --- | --- |
| 45 | Strain B::eri1Δ | T770.2h | CD429_i13_A | Input | 99.0 | 8010217 | N/A |  |
| 46 | Strain B::eri1Δ | T770.2h | CD238_i20_A | Input | 98.7 | 10293198 | N/A |  |
| 47 | Strain B::flag-sad-6 | T827.7h | CD528_i32_A | FLAG | 37.1 | 4506548 | CD527_i4_A | BI |
| 48 | Strain B::flag-sad-6 | T827.7h | CD763_i33_A | FLAG | 98.5 | 10738768 | CD672_i46_A | BI |
| 49 | Strain B::flag-sad-6 | T827.7h | CD764_i34_A | FLAG | 96.6 | 9150853 | CD672_i46_A | BI |
| 50 | Strain B::flag-sad-6 | T827.7h | CD765_i35_A | FLAG | 97.1 | 10576241 | CD677_i41_A | BI+Tc |
| 51 | Strain B::flag-sad-6 | T827.7h | CD766_i36_A | FLAG | 98.0 | 12321810 | CD677_i41_A | BI+Tc |
| 52 | Strain B::flag-sad-6 | T827.7h | CD675_i43_A | GFP | 97.9 | 10072417 | CD672_i46_A | BI |
| 53 | Strain B::flag-sad-6 | T827.7h | CD604_i39_A | GFP | 97.3 | 10537132 | CD601_i43_A | BI+Tc |
| 54 | Strain B::flag-sad-6 | T827.7h | CD680_i38_A | GFP | 98.4 | 10691467 | CD677_i41_A | BI+Tc |
| 55 | Strain B::flag-sad-6 | T827.7h | CD674_i44_A | H3K9me3 | 96.8 | 10377412 | CD672_i46_A | BI |
| 56 | Strain B::flag-sad-6 | T827.7h | CD799_i38_A | H3K9me3 | 91.3 | 5666258 | CD527_i4_A | BI |
| 57 | Strain B::flag-sad-6 | T827.7h | CD603_i40_A | H3K9me3 | 97.6 | 9444811 | CD601_i43_A | BI+Tc |
| 58 | Strain B::flag-sad-6 | T827.7h | CD679_i39_A | H3K9me3 | 97.3 | 10487142 | CD677_i41_A | BI+Tc |
| 59 | Strain B::flag-sad-6 | T827.7h | CD673_i45_A | hH3 | 98.6 | 10431332 | CD672_i46_A | BI |
| 60 | Strain B::flag-sad-6 | T827.7h | CD798_i37_A | hH3 | 98.8 | 7057779 | CD527_i4_A | BI |
| 61 | Strain B::flag-sad-6 | T827.7h | CD602_i41_A | hH3 | 98.8 | 10748400 | CD601_i43_A | BI+Tc |
| 62 | Strain B::flag-sad-6 | T827.7h | CD678_i40_A | hH3 | 98.8 | 10514385 | CD677_i41_A | BI+Tc |
| 63 | Strain B::flag-sad-6 | T827.7h | CD527_i4_A | Input | 98.9 | 10730984 | N/A | BI |
| 64 | Strain B::flag-sad-6 | T827.7h | CD672_i46_A | Input | 98.8 | 11006000 | N/A | BI |
| 65 | Strain B::flag-sad-6 | T827.7h | CD601_i43_A | Input | 98.9 | 10390350 | N/A | BI+Tc |
| 66 | Strain B::flag-sad-6 | T827.7h | CD677_i41_A | Input | 98.8 | 11240383 | N/A | BI+Tc |
| 67 | Strain B::hda1Δ | T714.8h | CD138_i34_A | H3K9me3 | 96.8 | 5320153 | CD136_i17_A |  |
| 68 | Strain B::hda1Δ | T714.8h | CD137_i19_A | hH3 | 99.1 | 7581904 | CD136_i17_A |  |
| 69 | Strain B::hda1Δ | T714.8h | CD136_i17_A | Input | 98.9 | 7478362 | N/A |  |
| 70 | Strain B::qde-1Δ | T710.2h | CD424_i1_A | GFP | 97.2 | 8583286 | CD421_i5_A |  |
| 71 | Strain B::qde-1Δ | T710.2h | CD435_i19_A | GFP | 93.1 | 7774689 | CD432_i16_A |  |
| 72 | Strain B::qde-1Δ | T710.2h | CD423_i7_A | H3K9me3 | 97.3 | 7300690 | CD421_i5_A |  |
| 73 | Strain B::qde-1Δ | T710.2h | CD434_i18_A | H3K9me3 | 97.2 | 8452778 | CD432_i16_A |  |
| 74 | Strain B::qde-1Δ | T710.2h | CD422_i6_A | hH3 | 99.0 | 8298992 | CD421_i5_A |  |
| 75 | Strain B::qde-1Δ | T710.2h | CD433_i17_A | hH3 | 98.9 | 9519976 | CD432_i16_A |  |
| 76 | Strain B::qde-1Δ | T710.2h | CD421_i5_A | Input | 99.0 | 8638738 | N/A |  |
| 77 | Strain B::qde-1Δ | T710.2h | CD432_i16_A | Input | 99.0 | 7704419 | N/A |  |
| 78 | Strain B::qde-2Δ | T711.21h | CD196_i32_A | H3K9me3 | 96.6 | 7747370 | CD194_i30_A |  |
| 79 | Strain B::qde-2Δ | T711.21h | CD438_i22_A | H3K9me3 | 97.0 | 8674141 | CD436_i20_A |  |
| 80 | Strain B::qde-2Δ | T711.21h | CD195_i31_A | hH3 | 98.5 | 8368280 | CD194_i30_A |  |
| 81 | Strain B::qde-2Δ | T711.21h | CD437_i21_A | hH3 | 69.3 | 6663839 | CD436_i20_A |  |
| 82 | Strain B::qde-2Δ | T711.21h | CD194_i30_A | Input | 98.5 | 8488470 | N/A |  |
| 83 | Strain B::qde-2Δ | T711.21h | CD436_i20_A | Input | 99.1 | 9493385 | N/A |  |
| 84 | Strain B::qde-3Δ | T712.2h | CD428_i12_A | GFP | 97.8 | 6900465 | CD425_i9_A |  |
| 85 | Strain B::qde-3Δ | T712.2h | CD445_i29_A | GFP | 98.6 | 6917632 | CD442_i26_A |  |
| 86 | Strain B::qde-3Δ | T712.2h | CD129_i44_A | H3K9me3 | 96.8 | 8243938 | CD127_i43_A |  |
| 87 | Strain B::qde-3Δ | T712.2h | CD444_i28_A | H3K9me3 | 97.2 | 8428380 | CD442_i26_A |  |
| 88 | Strain B::qde-3Δ | T712.2h | CD427_i11_A | H3K9me3 | 97.4 | 6846737 | CD425_i9_A |  |
| 89 | Strain B::qde-3Δ | T712.2h | CD128_i15_A | hH3 | 98.8 | 7664057 | CD127_i43_A |  |
| 90 | Strain B::qde-3Δ | T712.2h | CD426_i10_A | hH3 | 99.1 | 7094939 | CD425_i9_A |  |
| 91 | Strain B::qde-3Δ | T712.2h | CD443_i27_A | hH3 | 97.3 | 8516563 | CD442_i26_A |  |
| 92 | Strain B::qde-3Δ | T712.2h | CD127_i43_A | Input | 98.8 | 8261073 | N/A |  |
| 93 | Strain B::qde-3Δ | T712.2h | CD442_i26_A | Input | 99.1 | 6015297 | N/A |  |
| 94 | Strain B::qde-3Δ | T712.2h | CD425_i9_A | Input | 99.1 | 7512143 | N/A |  |

**Table S4. ChIP-seq libraries created and analyzed in this study (continued).**
|  |  |  |  |  |  |  |  |  |
| --- | --- | --- | --- | --- | --- | --- | --- | --- |
| 95 | Strain B::rad51Δ | T737.1h | CD147_i41_A | H3K9me3 | 97.0 | 8337203 | CD145_i39_A |  |
| 96 | Strain B::rad51Δ | T737.1h | CD404_i24_A | H3K9me3 | 97.2 | 10136955 | CD402_i22_A |  |
| 97 | Strain B::rad51Δ | T737.1h | CD403_i23_A | hH3 | 99.0 | 9258535 | CD402_i22_A |  |
| 98 | Strain B::rad51Δ | T737.1h | CD762_i32_A | hH3 | 98.4 | 8957878 | CD145_i39_A |  |
| 99 | Strain B::rad51Δ | T737.1h | CD145_i39_A | Input | 97.8 | 8378515 | N/A |  |
| 100 | Strain B::rad51Δ | T737.1h | CD402_i22_A | Input | 98.9 | 9409013 | N/A |  |
| 101 | Strain B::rad52Δ | T738.1h | CD186_i24_A | H3K9me3 | 96.5 | 9324486 | CD184_i22_A |  |
| 102 | Strain B::rad52Δ | T738.1h | CD407_i27_A | H3K9me3 | 97.1 | 9284593 | CD405_i25_A |  |
| 103 | Strain B::rad52Δ | T738.1h | CD185_i23_A | hH3 | 98.5 | 8814243 | CD184_i22_A |  |
| 104 | Strain B::rad52Δ | T738.1h | CD406_i26_A | hH3 | 99.0 | 8800775 | CD405_i25_A |  |
| 105 | Strain B::rad52Δ | T738.1h | CD184_i22_A | Input | 98.5 | 9206043 | N/A |  |
| 106 | Strain B::rad52Δ | T738.1h | CD405_i25_A | Input | 99.0 | 9393715 | N/A |  |
| 107 | Strain B::rtt109Δ | T786.3h | CD377_i48_A | H3K9me3 | 95.6 | 6482608 | CD375_i47_A |  |
| 108 | Strain B::rtt109Δ | T786.3h | CD448_i32_A | H3K9me3 | 97.2 | 7149877 | CD446_i30_A |  |
| 109 | Strain B::rtt109Δ | T786.3h | CD376_i29_A | hH3 | 98.3 | 8275565 | CD375_i47_A |  |
| 110 | Strain B::rtt109Δ | T786.3h | CD447_i31_A | hH3 | 99.0 | 7456714 | CD446_i30_A |  |
| 111 | Strain B::rtt109Δ | T786.3h | CD375_i47_A | Input | 98.7 | 7137760 | N/A |  |
| 112 | Strain B::rtt109Δ | T786.3h | CD446_i30_A | Input | 99.1 | 7804782 | N/A |  |
| 113 | Strain B::sad-6Δ | T707.51h | CD458_i42_A | GFP | 98.5 | 9891388 | CD454_i38_A | Tc |
| 114 | Strain B::sad-6Δ | T707.51h | CD632_i6_A | GFP | 98.8 | 8444453 | CD629_i9_A | Tc |
| 115 | Strain B::sad-6Δ | T707.51h | CD262_i43_A | GFP | 93.1 | 7346354 | CD259_i40_A |  |
| 116 | Strain B::sad-6Δ | T707.51h | CD401_i21_A | GFP | 98.2 | 8513589 | CD398_i18_A |  |
| 117 | Strain B::sad-6Δ | T707.51h | CD456_i40_A | H3K9me3 | 98.1 | 10193635 | CD454_i38_A | Tc |
| 118 | Strain B::sad-6Δ | T707.51h | CD631_i7_A | H3K9me3 | 97.4 | 9147582 | CD629_i9_A | Tc |
| 119 | Strain B::sad-6Δ | T707.51h | CD261_i42_A | H3K9me3 | 97.0 | 8832346 | CD259_i40_A |  |
| 120 | Strain B::sad-6Δ | T707.51h | CD400_i20_A | H3K9me3 | 97.0 | 7928262 | CD398_i18_A |  |
| 121 | Strain B::sad-6Δ | T707.51h | CD736_i31_A | hH2B | 99.0 | 8931671 | CD454_i38_A | Tc |
| 122 | Strain B::sad-6Δ | T707.51h | CD737_i32_A | hH2B | 98.9 | 7844732 | CD629_i9_A | Tc |
| 123 | Strain B::sad-6Δ | T707.51h | CD733_i28_A | hH2B | 98.9 | 9741353 | CD398_i18_A |  |
| 124 | Strain B::sad-6Δ | T707.51h | CD735_i30_A | hH2B | 98.8 | 9109414 | CD734_i29_A |  |
| 125 | Strain B::sad-6Δ | T707.51h | CD455_i39_A | hH3 | 99.0 | 13205725 | CD454_i38_A | Tc |
| 126 | Strain B::sad-6Δ | T707.51h | CD630_i8_A | hH3 | 98.8 | 9219792 | CD629_i9_A | Tc |
| 127 | Strain B::sad-6Δ | T707.51h | CD260_i41_A | hH3 | 98.5 | 7870943 | CD259_i40_A |  |
| 128 | Strain B::sad-6Δ | T707.51h | CD399_i19_A | hH3 | 98.9 | 9548072 | CD398_i18_A |  |
| 129 | Strain B::sad-6Δ | T707.51h | CD454_i38_A | Input | 99.2 | 11323585 | N/A | Tc |
| 130 | Strain B::sad-6Δ | T707.51h | CD629_i9_A | Input | 99.0 | 9775144 | N/A | Tc |
| 131 | Strain B::sad-6Δ | T707.51h | CD259_i40_A | Input | 98.8 | 12084203 | N/A |  |
| 132 | Strain B::sad-6Δ | T707.51h | CD398_i18_A | Input | 99.0 | 10727138 | N/A |  |
| 133 | Strain B::sad-6Δ | T707.51h | CD734_i29_A | Input | 98.8 | 8947082 | N/A |  |
| 134 | Strain C | C290.6 | CD69_i37_A | H3K9me3 | 98.4 | 7002290 | CD67_i35_A |  |
| 135 | Strain C | C290.6 | CD569_i45_A | H3K9me3 | 95.3 | 7793367 | CD567_i48_A |  |
| 136 | Strain C | C290.6 | CD68_i36_A | hH3 | 98.8 | 8373074 | CD67_i35_A |  |
| 137 | Strain C | C290.6 | CD568_i46_A | hH3 | 98.4 | 7145823 | CD567_i48_A |  |
| 138 | Strain C | C290.6 | CD67_i35_A | Input | 98.4 | 8426641 | N/A |  |
| 139 | Strain C | C290.6 | CD567_i48_A | Input | 98.5 | 8075101 | N/A |  |
| 140 | Strain C::lacO(d1); dim-5+ | T697.1 | CD61_i48_A | H3K9me3 | 97.8 | 5278648 | CD60_i47_A |  |
| 141 | Strain C::lacO(d1); dim-5+ | T697.1 | CD556_i36_A | H3K9me3 | 96.7 | 8153045 | CD555_i37_A |  |
| 142 | Strain C::lacO(d1); dim-5+ | T697.1 | CD60_i47_A | Input | 99.3 | 6408038 | N/A |  |
| 143 | Strain C::lacO(d1); dim-5+ | T697.1 | CD555_i37_A | Input | 99.0 | 9482890 | N/A |  |
| 144 | Strain C::lacO(d2); dim-5+ | T698.1 | CD66_i34_A | H3K9me3 | 88.5 | 7339481 | CD65_i33_A |  |

**Table S4. ChIP-seq libraries created and analyzed in this study (continued).**
|  |  |  |  |  |  |  |  |  |
| --- | --- | --- | --- | --- | --- | --- | --- | --- |
| 145 | Strain C::lacO(d2); dim-5+ | T698.1 | CD558_i34_A | H3K9me3 | 96.5 | 6864519 | CD557_i35_A |  |
| 146 | Strain C::lacO(d2); dim-5+ | T698.1 | CD65_i33_A | Input | 99.4 | 8631930 | N/A |  |
| 147 | Strain C::lacO(d2); dim-5+ | T698.1 | CD557_i35_A | Input | 98.9 | 8147564 | N/A |  |
| 148 | Strain C::dim-5+ | T663.2 | CD40_i27_A | H3K9me3 | 97.8 | 10427164 | CD38_i25_A |  |
| 149 | Strain C::dim-5+ | T663.2 | CD769_i38_A | H3K9me3 | 95.0 | 9142215 | CD294_i25_A |  |
| 150 | Strain C::dim-5+ | T663.2 | CD39_i26_A | hH3 | 99.0 | 7942226 | CD38_i25_A |  |
| 151 | Strain C::dim-5+ | T663.2 | CD768_i37_A | hH3 | 98.9 | 10418247 | CD294_i25_A |  |
| 152 | Strain C::dim-5+ | T663.2 | CD38_i25_A | Input | 99.2 | 6976064 | N/A |  |
| 153 | Strain C::dim-5+ | T663.2 | CD294_i25_A | Input | 98.4 | 9361089 | N/A |  |
| 154 | Strain C::sad-6Δ | T752.13h | CD830_i29_A | hH3 | 98.7 | 8808509 | CD829_i28_A |  |
| 155 | Strain C::sad-6Δ | T752.13h | CD829_i28_A | Input | 98.7 | 10053671 | N/A |  |
| 156 | Strain D::chapΔ | T690.1h | CD49_i36_A | H3K9me3 | 97.5 | 9337383 | CD47_i34_A |  |
| 157 | Strain D::chapΔ | T690.1h | CD579_i33_A | H3K9me3 | 97.2 | 8135547 | CD577_i35_A |  |
| 158 | Strain D::chapΔ | T690.1h | CD48_i35_A | hH3 | 99.2 | 8855879 | CD47_i34_A |  |
| 159 | Strain D::chapΔ | T690.1h | CD578_i34_A | hH3 | 98.6 | 7111554 | CD577_i35_A |  |
| 160 | Strain D::chapΔ | T690.1h | CD47_i34_A | Input | 99.1 | 6600607 | N/A |  |
| 161 | Strain D::chapΔ | T690.1h | CD577_i35_A | Input | 92.5 | 7972673 | N/A |  |
| 162 | Strain D::dim-7Δ | T688.2h | CD43_i30_A | H3K9me3 | 98.3 | 6777728 | CD41_i28_A |  |
| 163 | Strain D::dim-7Δ | T688.2h | CD573_i40_A | H3K9me3 | 94.2 | 8595604 | CD571_i43_A |  |
| 164 | Strain D::dim-7Δ | T688.2h | CD42_i29_A | hH3 | 99.2 | 11718495 | CD41_i28_A |  |
| 165 | Strain D::dim-7Δ | T688.2h | CD572_i41_A | hH3 | 98.5 | 7047100 | CD571_i43_A |  |
| 166 | Strain D::dim-7Δ | T688.2h | CD41_i28_A | Input | 99.1 | 8792743 | N/A |  |
| 167 | Strain D::dim-7Δ | T688.2h | CD571_i43_A | Input | 98.6 | 8428865 | N/A |  |
| 168 | Strain D::dim-9Δ | T689.2h | CD46_i33_A | H3K9me3 | 98.4 | 8255276 | CD44_i31_A |  |
| 169 | Strain D::dim-9Δ | T689.2h | CD576_i36_A | H3K9me3 | 96.9 | 9271303 | CD574_i39_A |  |
| 170 | Strain D::dim-9Δ | T689.2h | CD45_i32_A | hH3 | 99.2 | 9513255 | CD44_i31_A |  |
| 171 | Strain D::dim-9Δ | T689.2h | CD575_i38_A | hH3 | 98.8 | 9986908 | CD574_i39_A |  |
| 172 | Strain D::dim-9Δ | T689.2h | CD44_i31_A | Input | 99.2 | 6616746 | N/A |  |
| 173 | Strain D::dim-9Δ | T689.2h | CD574_i39_A | Input | 98.5 | 10005228 | N/A |  |
| 174 | Strain D::hda1Δ | T743.3h | CD135_i48_A | H3K9me3 | 97.1 | 6833539 | CD133_i47_A |  |
| 175 | Strain D::hda1Δ | T743.3h | CD134_i18_A | hH3 | 98.9 | 8831960 | CD133_i47_A |  |
| 176 | Strain D::hda1Δ | T743.3h | CD133_i47_A | Input | 98.9 | 6330925 | N/A |  |
| 177 | Strain E | T601.2h | CD848_i36_A | H3K9me3 | 93.7 | 8643593 | CD845_i33_A | "maintained" |
| 178 | Strain E | T601.2h | CD844_i42_A | H3K9me3 | 96.6 | 7761265 | CD843_i41_A | "released" |
| 179 | Strain E | T601.2h | CD852_i4_A | H3K9me3 | 92.3 | 8634000 | CD849_i1_A | "released" |
| 180 | Strain E | T601.2h | CD655_i37_A | H3K9me3 | 98.1 | 8476509 | CD653_i35_A | no-N |
| 181 | Strain E | T601.2h | CD750_i25_A | H3K9me3 | 97.4 | 8908047 | CD723_i18_A | no-N |
| 182 | Strain E | T601.2h | CD75_i43_A | H3K9me3 | 97.5 | 7793876 | CD73_i41_A |  |
| 183 | Strain E | T601.2h | CD652_i34_A | H3K9me3 | 98.5 | 8145259 | CD650_i32_A |  |
| 184 | Strain E | T601.2h | CD741_i36_A | hH2B | 98.8 | 10530685 | CD723_i18_A | no-N |
| 185 | Strain E | T601.2h | CD740_i35_A | hH2B | 98.9 | 8585096 | CD697_i24_A | no-N |
| 186 | Strain E | T601.2h | CD738_i33_A | hH2B | 98.8 | 9637627 | CD650_i32_A |  |
| 187 | Strain E | T601.2h | CD739_i34_A | hH2B | 99.0 | 8700628 | CD721_i16_A |  |
| 188 | Strain E | T601.2h | CD654_i36_A | hH3 | 96.3 | 7792030 | CD653_i35_A | no-N |
| 189 | Strain E | T601.2h | CD698_i23_A | hH3 | 98.6 | 8852422 | CD697_i24_A | no-N |
| 190 | Strain E | T601.2h | CD74_i42_A | hH3 | 99.1 | 8816753 | CD73_i41_A |  |
| 191 | Strain E | T601.2h | CD651_i33_A | hH3 | 99.1 | 8817367 | CD650_i32_A |  |
| 192 | Strain E | T601.2h | CD845_i33_A | Input | 99.1 | 9926094 | N/A | "maintained" |
| 193 | Strain E | T601.2h | CD843_i41_A | Input | 98.9 | 9427679 | N/A | "released" |
| 194 | Strain E | T601.2h | CD849_i1_A | Input | 99.0 | 9054557 | N/A | "released" |

**Table S4. ChIP-seq libraries created and analyzed in this study (continued).**
|  |  |  |  |  |  |  |  |  |
| --- | --- | --- | --- | --- | --- | --- | --- | --- |
| 195 | Strain E | T601.2h | CD723_i18_A | Input | 99.0 | 9226469 | N/A | no-N |
| 196 | Strain E | T601.2h | CD697_i24_A | Input | 99.0 | 8445763 | N/A | no-N |
| 197 | Strain E | T601.2h | CD653_i35_A | Input | 99.0 | 9196543 | N/A | no-N |
| 198 | Strain E | T601.2h | CD650_i32_A | Input | 98.6 | 9355163 | N/A |  |
| 199 | Strain E | T601.2h | CD721_i16_A | Input | 98.9 | 11369712 | N/A |  |
| 200 | Strain E | T601.2h | CD73_i41_A | Input | 98.0 | 9209709 | N/A |  |
| 201 | Strain E::tetR-gfp | T753.1 | CD190_i28_A | H3K9me3 | 96.8 | 8798934 | CD188_i26_A |  |
| 202 | Strain E::tetR-gfp | T753.1 | CD614_i30_A | H3K9me3 | 97.4 | 7039957 | CD612_i32_A |  |
| 203 | Strain E::tetR-gfp | T753.1 | CD189_i27_A | hH3 | 98.5 | 8198700 | CD188_i26_A |  |
| 204 | Strain E::tetR-gfp | T753.1 | CD613_i31_A | hH3 | 99.0 | 9640803 | CD612_i32_A |  |
| 205 | Strain E::tetR-gfp | T753.1 | CD188_i26_A | Input | 98.6 | 7988786 | N/A |  |
| 206 | Strain E::tetR-gfp | T753.1 | CD612_i32_A | Input | 98.9 | 11275351 | N/A |  |
| 207 | Strain E::nit-2Δ | T851.1h | CD667_i48_A | H3K9me3 | 98.4 | 8032158 | CD665_i46_A | no-N |
| 208 | Strain E::nit-2Δ | T851.1h | CD690_i28_A | H3K9me3 | 96.3 | 9369072 | CD688_i30_A | no-N |
| 209 | Strain E::nit-2Δ | T851.1h | CD666_i47_A | hH3 | 98.4 | 9158422 | CD665_i46_A | no-N |
| 210 | Strain E::nit-2Δ | T851.1h | CD689_i29_A | hH3 | 98.4 | 9083438 | CD688_i30_A | no-N |
| 211 | Strain E::nit-2Δ | T851.1h | CD665_i46_A | Input | 99.1 | 10457276 | N/A | no-N |
| 212 | Strain E::nit-2Δ | T851.1h | CD688_i30_A | Input | 98.9 | 9823600 | N/A | no-N |
| 213 | Strain E::qde-1Δ | T839.10h | CD608_i35_A | H3K9me3 | 88.5 | 10022605 | CD606_i36_A | no-N |
| 214 | Strain E::qde-1Δ | T839.10h | CD711_i6_A | H3K9me3 | 98.2 | 6636860 | CD709_i4_A | no-N |
| 215 | Strain E::qde-1Δ | T839.10h | CD607_i27_A | hH3 | 98.6 | 9631010 | CD606_i36_A | no-N |
| 216 | Strain E::qde-1Δ | T839.10h | CD710_i5_A | hH3 | 98.8 | 8712589 | CD709_i4_A | no-N |
| 217 | Strain E::qde-1Δ | T839.10h | CD606_i36_A | Input | 98.9 | 12868225 | N/A | no-N |
| 218 | Strain E::qde-1Δ | T839.10h | CD709_i4_A | Input | 99.2 | 8851401 | N/A | no-N |
| 219 | Strain E::qde-3Δ | T840.10h | CD611_i33_A | H3K9me3 | 97.7 | 10392870 | CD609_i34_A | no-N |
| 220 | Strain E::qde-3Δ | T840.10h | CD752_i26_A | H3K9me3 | 97.7 | 9109045 | CD715_i10_A | no-N |
| 221 | Strain E::qde-3Δ | T840.10h | CD610_i26_A | hH3 | 86.9 | 7683121 | CD609_i34_A | no-N |
| 222 | Strain E::qde-3Δ | T840.10h | CD716_i11_A | hH3 | 99.0 | 7553969 | CD715_i10_A | no-N |
| 223 | Strain E::qde-3Δ | T840.10h | CD715_i10_A | Input | 99.2 | 9094349 | N/A | no-N |
| 224 | Strain E::qde-3Δ | T840.10h | CD609_i34_A | Input | 99.1 | 10277658 | N/A | no-N |
| 225 | Strain E::rad51Δ | T847.3h | CD708_i3_A | H3K9me3 | 77.4 | 5436037 | CD706_i1_A | no-N |
| 226 | Strain E::rad51Δ | T847.3h | CD754_i27_A | H3K9me3 | 97.8 | 8943634 | CD700_i21_A | no-N |
| 227 | Strain E::rad51Δ | T847.3h | CD701_i20_A | hH3 | 99.0 | 9300497 | CD700_i21_A | no-N |
| 228 | Strain E::rad51Δ | T847.3h | CD707_i2_A | hH3 | 97.7 | 8549931 | CD706_i1_A | no-N |
| 229 | Strain E::rad51Δ | T847.3h | CD700_i21_A | Input | 99.0 | 10417937 | N/A | no-N |
| 230 | Strain E::rad51Δ | T847.3h | CD706_i1_A | Input | 99.1 | 7828349 | N/A | no-N |
| 231 | Strain E::rad52Δ | T848.1h | CD714_i9_A | H3K9me3 | 97.0 | 7071655 | CD712_i7_A | no-N |
| 232 | Strain E::rad52Δ | T848.1h | CD756_i28_A | H3K9me3 | 97.7 | 9872070 | CD703_i18_A | no-N |
| 233 | Strain E::rad52Δ | T848.1h | CD704_i17_A | hH3 | 98.5 | 9440003 | CD703_i18_A | no-N |
| 234 | Strain E::rad52Δ | T848.1h | CD713_i8_A | hH3 | 98.8 | 7385596 | CD712_i7_A | no-N |
| 235 | Strain E::rad52Δ | T848.1h | CD703_i18_A | Input | 99.0 | 8137387 | N/A | no-N |
| 236 | Strain E::rad52Δ | T848.1h | CD712_i7_A | Input | 99.0 | 8000728 | N/A | no-N |
| 237 | Strain E::sad-6Δ | T774.4h | CD658_i40_A | H3K9me3 | 97.7 | 9927567 | CD656_i38_A | no-N |
| 238 | Strain E::sad-6Δ | T774.4h | CD687_i31_A | H3K9me3 | 96.5 | 9708825 | CD685_i33_A | no-N |
| 239 | Strain E::sad-6Δ | T774.4h | CD745_i40_A | hH2B | 98.9 | 7706122 | CD685_i33_A | no-N |
| 240 | Strain E::sad-6Δ | T774.4h | CD746_i41_A | hH2B | 98.7 | 7986318 | CD727_i22_A | no-N |
| 241 | Strain E::sad-6Δ | T774.4h | CD742_i37_A | hH2B | 99.0 | 10163595 | CD725_i20_A |  |
| 242 | Strain E::sad-6Δ | T774.4h | CD744_i39_A | hH2B | 98.8 | 9149919 | CD743_i38_A |  |
| 243 | Strain E::sad-6Δ | T774.4h | CD657_i39_A | hH3 | 98.5 | 9787164 | CD656_i38_A | no-N |
| 244 | Strain E::sad-6Δ | T774.4h | CD686_i32_A | hH3 | 98.5 | 9187849 | CD685_i33_A | no-N |

**Table S4. ChIP-seq libraries created and analyzed in this study (continued).**
|  |  |  |  |  |  |  |  |  |
| --- | --- | --- | --- | --- | --- | --- | --- | --- |
| 245 | Strain E::sad-6Δ | T774.4h | CD685_i33_A | Input | 98.9 | 8575543 | N/A | no-N |
| 246 | Strain E::sad-6Δ | T774.4h | CD656_i38_A | Input | 99.0 | 9538401 | N/A | no-N |
| 247 | Strain E::sad-6Δ | T774.4h | CD727_i22_A | Input | 99.0 | 8307630 | N/A | no-N |
| 248 | Strain E::sad-6Δ | T774.4h | CD725_i20_A | Input | 98.9 | 9546376 | N/A |  |
| 249 | Strain E::sad-6Δ | T774.4h | CD743_i38_A | Input | 98.4 | 9014884 | N/A |  |
| 250 | Strain F::lacO(i1); dim-5+ | T764.1 | CD202_i38_A | H3K9me3 | 96.9 | 7305893 | CD201_i37_A |  |
| 251 | Strain F::lacO(i1); dim-5+ | T764.1 | CD562_i30_A | H3K9me3 | 96.8 | 9060889 | CD561_i31_A |  |
| 252 | Strain F::lacO(i1); dim-5+ | T764.1 | CD201_i37_A | Input | 98.5 | 6529896 | N/A |  |
| 253 | Strain F::lacO(i1); dim-5+ | T764.1 | CD561_i31_A | Input | 98.8 | 10258869 | N/A |  |
| 254 | Strain F::lacO(i2); dim-5+ | T766.1 | CD204_i40_A | H3K9me3 | 96.1 | 5680629 | CD203_i39_A |  |
| 255 | Strain F::lacO(i2); dim-5+ | T766.1 | CD564_i28_A | H3K9me3 | 96.6 | 8147239 | CD563_i29_A |  |
| 256 | Strain F::lacO(i2); dim-5+ | T766.1 | CD203_i39_A | Input | 98.4 | 8587633 | N/A |  |
| 257 | Strain F::lacO(i2); dim-5+ | T766.1 | CD563_i29_A | Input | 98.5 | 8521230 | N/A |  |
| 258 | Strain F::lacO(i3); dim-5+ | T776.1 | CD206_i42_A | H3K9me3 | 96.7 | 6528076 | CD205_i41_A |  |
| 259 | Strain F::lacO(i3); dim-5+ | T776.1 | CD566_i42_A | H3K9me3 | 97.2 | 7423559 | CD565_i47_A |  |
| 260 | Strain F::lacO(i3); dim-5+ | T776.1 | CD205_i41_A | Input | 98.6 | 5535835 | N/A |  |
| 261 | Strain F::lacO(i3); dim-5+ | T776.1 | CD565_i47_A | Input | 98.9 | 8428602 | N/A |  |

**Table S5.** MNase-seq libraries created and analyzed in this study.

| # | Genetic condition | Strain ID | Library | Alignment rate (%) | Aligned reads * | Additional conditions |
| --- | --- | --- | --- | --- | --- | --- |
| 1 | Strain A | T647.1h | CD1072_i0_A | 99.3 | 26796362 |  |
| 2 | Strain B | T658.1 | CD530_i12_A | 99.1 | 10769647 |  |
| 3 | Strain B::sad-6Δ | T707.51h | CD533_i15_A | 99.0 | 12371585 |  |
| 4 | Strain C | C290.6 | CD290_i21_A | 98.5 | 11797272 |  |
| 5 | Strain C::sad-6Δ | T752.13h | CD832_i30_A | 99.2 | 12764340 |  |
| 6 | Strain E | T601.2h | CD722_i17_A | 99.1 | 11456746 |  |
| 7 | Strain E | T601.2h | CD587_i15_A | 99.2 | 15232062 | no-N |
| 8 | Strain E::sad-6Δ | T774.4h | CD589_i14_A | 99.3 | 14815243 | no-N |

\* Number of aligned reads is reported for the nuclear genome (corresponding to the interval 64841-41123470 in *mNC12.lacO-tetO*)

**Table S6.** sRNA-seq libraries created and analyzed in this study.

| # | Genetic condition | Strain ID | Library | Alignment rate (%) | Mitochondrial reads | Ribosomal reads | Core nuclear reads | Additional conditions |
| --- | --- | --- | --- | --- | --- | --- | --- | --- |
| 1 | Strain A | T647.1h | VR18_i16_A | 91.5 | 89140 | 6334645 | 768640 |  |
| 2 | Strain A | T647.1h | VR167_i2_A | 94.5 | 273129 | 4821765 | 1112917 |  |
| 3 | Strain B | T658.1 | VR13_i11_A | 94.7 | 72864 | 3349707 | 669524 |  |
| 4 | Strain B | T658.1 | VR200_i19_A | 95.1 | 354937 | 6246084 | 2046200 |  |
| 5 | Strain B | T658.1 | VR259_i47_A | 92.6 | 330465 | 4767444 | 1953046 | BI |
| 6 | Strain B | T658.1 | VR274_i37_A | 95.3 | 397237 | 6514549 | 2298582 | BI |
| 7 | Strain B | T658.1 | VR48_i25_A | 93.6 | 1591285 | 3207673 | 923786 | HU |
| 8 | Strain B | T658.1 | VR202_i21_A | 94.3 | 1523622 | 4343824 | 2032401 | HU |
| 9 | Strain B | T658.1 | VR201_i20_A | 95.9 | 283061 | 6067766 | 1137260 | Tc |
| 10 | Strain B | T658.1 | VR232_i43_A | 95.0 | 100545 | 7477216 | 1011309 | Tc |
| 11 | Strain B::chd1Δ | T709.3h | VR181_i9_A | 95.1 | 365062 | 5327883 | 2700055 |  |
| 12 | Strain B::chd1Δ | T709.3h | VR234_i45_A | 95.0 | 228614 | 4714202 | 1942622 |  |
| 13 | Strain B::eri1Δ | T770.2h | VR156_i18_A | 93.7 | 208510 | 3976531 | 2827441 |  |
| 14 | Strain B::eri1Δ | T770.2h | VR226_i43_A | 94.6 | 168717 | 5107627 | 2397763 |  |
| 15 | Strain B::flag-sad-6 | T827.7h | VR275_i38_A | 95.2 | 362259 | 6686193 | 1841027 | BI |
| 16 | Strain B::flag-sad-6 | T827.7h | VR328_i0_A | 86.5 | 140179 | 4821635 | 969352 | BI |
| 17 | Strain B::hda1Δ | T714.8h | VR67_i20_A | 96.4 | 117222 | 1653395 | 4463222 |  |
| 18 | Strain B::hda1Δ | T714.8h | VR187_i5_A | 93.3 | 180826 | 2141171 | 5662706 |  |
| 19 | Strain B::qde-1Δ | T710.2h | VR51_i28_A | 94.1 | 199541 | 5057507 | 1055664 |  |
| 20 | Strain B::qde-1Δ | T710.2h | VR158_i20_A | 93.8 | 337753 | 5208452 | 1259318 |  |
| 21 | Strain B::qde-2Δ | T711.21h | VR159_i21_A | 91.0 | 176974 | 3485602 | 2274100 |  |
| 22 | Strain B::qde-2Δ | T711.21h | VR206_i25_A | 90.5 | 352184 | 6498566 | 3417528 |  |
| 23 | Strain B::qde-3Δ | T712.2h | VR75_i33_A | 96.0 | 175742 | 4122871 | 622478 |  |
| 24 | Strain B::qde-3Δ | T712.2h | VR207_i26_A | 90.3 | 177939 | 5426733 | 786112 |  |
| 25 | Strain B::rad51Δ | T737.1h | VR182_i46_A | 94.9 | 272390 | 5804726 | 3582359 |  |
| 26 | Strain B::rad51Δ | T737.1h | VR219_i36_A | 93.3 | 285179 | 7066045 | 2967365 |  |
| 27 | Strain B::rad52Δ | T738.1h | VR178_i42_A | 95.2 | 234040 | 3694120 | 7134007 |  |
| 28 | Strain B::rad52Δ | T738.1h | VR210_i29_A | 94.0 | 213286 | 3682292 | 4382441 |  |
| 29 | Strain B::rtt109Δ | T786.3h | VR213_i32_A | 94.0 | 127733 | 6865730 | 1343029 |  |
| 30 | Strain B::rtt109Δ | T786.3h | VR221_i38_A | 94.4 | 171898 | 6281191 | 1998765 |  |
| 31 | Strain B::sad-6Δ | T707.51h | VR157_i19_A | 92.4 | 384973 | 5419815 | 1623912 |  |
| 32 | Strain B::sad-6Δ | T707.51h | VR204_i23_A | 95.4 | 305828 | 5999418 | 905752 |  |
| 33 | Strain B::top3Δ | T812.11h | VR242_i42_A | 95.8 | 175430 | 3631633 | 4186739 |  |
| 34 | Strain B::top3Δ | T812.11h | VR246_i46_A | 94.8 | 122062 | 5173798 | 2023411 |  |
| 35 | Strain C | C290.6 | VR139_i1_A | 94.0 | 270021 | 5810093 | 2915944 |  |
| 36 | Strain C | C290.6 | VR195_i14_A | 90.4 | 192356 | 3984691 | 2252637 |  |
| 37 | Strain C | C290.6 | VR15_i13_A | 88.9 | 598556 | 2738413 | 756356 | HU |
| 38 | Strain C | C290.6 | VR199_i18_A | 93.8 | 296056 | 5583631 | 1021777 | HU |
| 39 | Strain C::lacO(d1) | C290.6.C4 | VR4_i3_A | 95.4 | 101949 | 3745549 | 961248 |  |
| 40 | Strain C::lacO(d1) | C290.6.C4 | VR264_i27_A | 96.5 | 248698 | 6752046 | 2180782 |  |
| 41 | Strain C::lacO(d2) | C290.6.C6 | VR5_i28_A | 92.3 | 124222 | 3910687 | 1078488 |  |
| 42 | Strain C::lacO(d2) | C290.6.C6 | VR265_i28_A | 93.8 | 334516 | 6952769 | 3550683 |  |
| 43 | Strain C::dim-5+ | T663.2 | VR1_i27_A | 96.3 | 176973 | 5096301 | 894206 |  |
| 44 | Strain C::dim-5+ | T663.2 | VR140_i2_A | 95.2 | 323739 | 9390098 | 1655988 |  |

**Table S6. sRNA-seq libraries created and analyzed in this study (continued).**
|  |  |  |  |  |  |  |  |  |
| --- | --- | --- | --- | --- | --- | --- | --- | --- |
| 45 | Strain C::qde-1Δ | T739.4h | VR52_i29_A | 96.3 | 167825 | 5736656 | 849642 |  |
| 46 | Strain C::qde-1Δ | T739.4h | VR170_i34_A | 94.6 | 63998 | 8986893 | 962974 |  |
| 47 | Strain C::qde-1Δ | T739.4h | VR368_i2_A | 97.9 | 254400 | 22662127 | 2471385 |  |
| 48 | Strain C::rad51Δ | T678.5h | VR27_i23_A | 95.2 | 186305 | 6174654 | 2362345 |  |
| 49 | Strain C::rad51Δ | T678.5h | VR41_i18_A | 90.0 | 62074 | 2358548 | 1006793 |  |
| 50 | Strain C::rad51Δ | T678.5h | VR177_i41_A | 94.3 | 215374 | 4479746 | 5634639 |  |
| 51 | Strain C::sad-6Δ | T752.13h | VR255_i39_A | 96.1 | 153384 | 9522494 | 1376775 |  |
| 52 | Strain C::sad-6Δ | T752.13h | VR260_i48_A | 95.9 | 143268 | 7577198 | 1006200 |  |
| 53 | Strain D::chapΔ | T690.1h | VR186_i26_A | 91.5 | 830143 | 4924184 | 2136729 |  |
| 54 | Strain D::chapΔ | T690.1h | VR198_i17_A | 94.7 | 166768 | 5612364 | 1034516 |  |
| 55 | Strain D::dim-7Δ | T688.2h | VR6_i4_A | 96.5 | 161363 | 5416165 | 1501359 |  |
| 56 | Strain D::dim-7Δ | T688.2h | VR143_i5_A | 94.2 | 185216 | 7371866 | 2690269 |  |
| 57 | Strain D::dim-9Δ | T689.2h | VR7_i5_A | 95.5 | 119282 | 3553738 | 1260019 |  |
| 58 | Strain D::dim-9Δ | T689.2h | VR266_i29_A | 95.3 | 225356 | 7131021 | 2355417 |  |
| 59 | Strain D::hda1Δ | T743.3h | VR62_i13_A | 94.1 | 85391 | 1872060 | 2123860 |  |
| 60 | Strain D::hda1Δ | T743.3h | VR179_i43_A | 93.4 | 209183 | 4538707 | 4005878 |  |
| 61 | Strain E | T601.2h | VR166_i30_A | 94.5 | 435793 | 6926994 | 1351635 |  |
| 62 | Strain E | T601.2h | VR347_i0_A | 93.0 | 304726 | 12151367 | 1691174 |  |
| 63 | Strain E | T601.2h | VR304_i23_A | 95.2 | 312596 | 7185662 | 2676606 | no-N |
| 64 | Strain E | T601.2h | VR308_i1_A | 94.5 | 384311 | 6889801 | 1990153 | no-N |
| 65 | Strain E | T601.2h | VR280_i43_A | 94.3 | 176832 | 4934667 | 894476 | N+ control |
| 66 | Strain E | T601.2h | VR350_i48_A | 93.8 | 46571 | 4022817 | 559369 | N+ control |
| 67 | Strain E | T601.2h | VR353_i46_A | 94.2 | 134854 | 5476783 | 813063 | N+ control |
| 68 | Strain E | T601.2h | VR314_i7_A | 94.3 | 678899 | 9254058 | 1398482 | "released" |
| 69 | Strain E | T601.2h | VR315_i8_A | 94.0 | 330082 | 7129312 | 1850101 | "maintained" |
| 70 | Strain E::tetR-gfp | T753.1 | VR180_i44_A | 89.2 | 231596 | 8574884 | 2001104 |  |
| 71 | Strain E::tetR-gfp | T753.1 | VR208_i27_A | 93.8 | 256389 | 6289769 | 2410247 |  |
| 72 | Strain E::nit-2Δ | T851.1h | VR329_i0_A | 93.2 | 290039 | 8427720 | 1701623 | no-N |
| 73 | Strain E::nit-2Δ | T851.1h | VR330_i0_A | 94.5 | 250838 | 8409414 | 1660895 | no-N |
| 74 | Strain E::nit-2Δ | T851.1h | VR344_i5_A | 95.4 | 194513 | 6910524 | 1270996 | no-N |
| 75 | Strain E::qde-1Δ | T839.10h | VR312_i0_A | 95.4 | 889847 | 17261386 | 2543135 | no-N |
| 76 | Strain E::qde-1Δ | T839.10h | VR316_i0_A | 95.1 | 521087 | 11517703 | 1726187 | no-N |
| 77 | Strain E::qde-3Δ | T840.10h | VR313_i0_A | 95.1 | 917304 | 16945619 | 2790338 | no-N |
| 78 | Strain E::qde-3Δ | T840.10h | VR317_i0_A | 95.2 | 746121 | 12363669 | 2086566 | no-N |
| 79 | Strain E::rad51Δ | T847.3h | VR331_i3_B | 94.2 | 181149 | 3410221 | 2840787 | no-N |
| 80 | Strain E::rad51Δ | T847.3h | VR332_i4_C | 94.3 | 245869 | 5105236 | 4387030 | no-N |
| 81 | Strain E::rad52Δ | T848.1h | VR333_i5_B | 92.9 | 129785 | 3427957 | 2487608 | no-N |
| 82 | Strain E::rad52Δ | T848.1h | VR334_i6_B | 90.7 | 103778 | 3446758 | 1789739 | no-N |
| 83 | Strain E::sad-6Δ | T774.4h | VR302_i0_A | 94.6 | 397546 | 10630045 | 1666933 | no-N |
| 84 | Strain E::sad-6Δ | T774.4h | VR348_i0_A | 95.0 | 431348 | 12883573 | 2143730 | no-N |
| 85 | Strain F::lacO(i1) | T724.2h | VR76_i4_A | 91.1 | 121473 | 3805143 | 899079 |  |
| 86 | Strain F::lacO(i1) | T724.2h | VR189_i7_A | 90.8 | 272686 | 3757672 | 2319565 |  |
| 87 | Strain F::lacO(i2) | T726.1h | VR80_i8_A | 93.5 | 132936 | 3561318 | 1139354 |  |
| 88 | Strain F::lacO(i2) | T726.1h | VR191_i10_A | 91.8 | 467307 | 4934028 | 2998980 |  |
| 89 | Strain F::lacO(i3) | T728.1h | VR81_i25_A | 85.6 | 62911 | 3848337 | 583177 |  |

In Tables S4-S6, the following notations of the culturing conditions were used: ‘Tc’ - growing cultures were supplemented with 100 μg/μl of tetracycline;

’Bl’ - growing cultures were supplemented with 0.1 μg/μl of blasticidin S;

’HU’ - pre-grown cultures were incubated for 24 hours in the standard medium supplemented with 0.1M HU;

’no-N’ - cultures were shifted to the nitrogen-free medium without HU block;

’N+ control’ - cultures were processed analogously to ‘no-N’ but shifted to the standard (N+) medium instead;

“maintained” - cultures were treated with HU and shifted to the nitrogen-free medium containing HU;

“released” - cultures were treated with HU and shifted to the nitrogen-free medium without HU.

### SI DATA FILES

Available at https://figshare.com/s/68987d7763146c2a4e2c; DOI: 10.6084/m9.figshare.24943173

**Data File 1. Maps of plasmids used in this study.**

Provided as plain-text files in EMBL format, compressed with TAR/GZIP.

**Data File 2. Custom genome references used in this study.**

Provided as plain-text files in EMBL format, compressed with TAR/GZIP. The following genome references are included: *mNC12.lacO-tetO*, *mNC12.tetO*, and *mito_ribo*.

**Data File 3. 500-bp tiles representing reference sRNA-seq loci.**

Provided as a plain-text file in CSV format, compressed with TAR/GZIP. Tile coordinates correspond to the genome reference *mNC12.lacO-tetO*.

